# Differential developmental blueprints of organ-intrinsic nervous systems

**DOI:** 10.1101/2023.12.12.571306

**Authors:** I-Uen Hsu, Yingxin Lin, Yunshan Guo, Qian J. Xu, Yuancheng Shao, Ruiqi L. Wang, Dominic Yin, Jia Zhao, Lawrence H. Young, Hongyu Zhao, Le Zhang, Rui B. Chang

## Abstract

The organ-intrinsic nervous system is a major interface between visceral organs and the brain, mediating important sensory and regulatory functions in the body-brain axis and serving as critical local processors for organ homeostasis. Molecularly, anatomically, and functionally, organ-intrinsic neurons are highly specialized for their host organs. However, the underlying mechanism that drives this specialization is largely unknown. Here, we describe the differential strategies utilized to achieve organ-specific organization between the enteric nervous system (ENS)^1^ and the intrinsic cardiac nervous system (ICNS)^2^, a neuronal network essential for heart performance but poorly characterized. Integrating high-resolution whole-embryo imaging, single-cell genomics, spatial transcriptomics, proteomics, and bioinformatics, we uncover that unlike the ENS which is highly mobile and colonizes the entire gastrointestinal (GI) tract, the ICNS uses a rich set of extracellular matrix (ECM) genes that match with surrounding heart cells and an intermediate dedicated neuronal progenitor state to stabilize itself for a ‘beads-on-the-necklace’ organization on heart atria. While ICNS- and ENS-precursors are genetically similar, their differentiation paths are influenced by their host-organs, leading to distinct mature neuron types. Co-culturing ENS-precursors with heart cells shifts their identity towards the ICNS and induces the expression of heart-matching ECM genes. Our cross-organ study thus reveals fundamental principles for the maturation and specialization of organ-intrinsic neurons.

## Main

Many visceral organs have their own intrinsic nervous system, including the gut (enteric nervous system, ENS, ‘second brain’ on the gut^1^), heart (the intrinsic cardiac nervous system, ICNS, ‘little brain’ on the heart^2^), pancreas (intrapancreatic neurons^3,4^), and lung (intrapulmonary neurons^5,6^, Extended Data Fig. 1). These organ-intrinsic neural networks directly interact with various types of cells within their host organs, forming an immediate interface between visceral organs and the nervous system. Previous studies have shown that organ-intrinsic neurons are essential for organ functions. For example, the ENS regulates key digestive and immune functions such as gut motility, immunity, intestinal barrier integrity, and microbiome homeostasis, and is also critically involved in many psychological and neurological disorders^7,8^. Likewise, the ICNS plays a critical role in maintaining cardiac hemodynamics^2,9–12^ and intrapancreatic neurons potently regulate pancreatic endocrine secretion^3^. Therefore, organ-intrinsic neurons represent an essential component of the body-brain axis and are critical for organs’ pathophysiology.

Although organ-intrinsic neurons across different visceral organs share the same developmental origin of the neural crest^13^ and express a similar set of neurochemicals, increasing evidence suggests that they are highly specialized in organization patterns and molecular compositions for their host organs. For example, the gut is densely colonized by millions of enteric neurons^7^ but neurons on the heart, lung, and pancreas are much fewer in number and spatially limited to only a few locations^3,5,12^ (Extended Data Fig. 1). Within the ENS, neurons also display region-specific cytoarchitecture and subpopulations along the gastrointestinal (GI) tract^14^. These differences in cellular organizations and molecular architectures indicate that organ-intrinsic neuronal networks are shaped by their unique organ environment to achieve organ-specific functions. However, our knowledge about organ-intrinsic neurons other than the ENS is very limited, so that how they become specialized for their host organs remains mysterious.

Here we focus on the ICNS, an essential heart-brain interface that controls key cardiac functions and serves as a therapeutic target for debilitating cardiac diseases^15–22^. Using a collection of genetic and imaging tools, we for the first time uncover critical anatomical and molecular steps during the development of the ICNS. Through comparisons with the ENS, we further elucidate how organ environment influences the maturation and specialization of organ-intrinsic neurons. Our cross-organ study thus reveals differential blueprints for the formation of the organ-intrinsic neural network.

### A ‘beads-on-the-necklace’ architecture of the ICNS generated from cardiac cell growth

In contrast to the ENS, only ∼1,500 intrinsic cardiac neurons (ICNS-neurons) are distributed in a few discrete ganglia at stereotypical locations on the atrial surface of adult mouse heart^12^ (Fig. 1a, Extended Data Fig. 2a). We first performed lineage tracing to precisely define ICNS-neuron origin using a collection of Cre and Flpo mouse lines. ICNS-neurons labelled using a pan-neuronal marker mouse line Baf53b-Cre^23^ actively express PHOX2B (a pan-organ-intrinsic neuron master regulator^24^, Fig. 1b-c) and are effectively labelled in Wnt1-Cre; lox-Sun1-sfGFP^25^ (Wnt1^Sun1-GFP^), Sox10^Sun1-GFP^, and Phox2b-Flpo; frt-tdTomato (Phox2b^tdT^) mice (Extended Data Fig. 2), indicating that ICNS-neurons are developed from neural-crest derived cells^26^. To visualize ICNS migration and ganglion formation, we performed 3D imaging of optically cleared C57BL/6J mouse embryos against PHOX2B. Our data show that ICNS cells first land on the heart at embryonic day (E) 12.5 via vagal cardiac branches, as reported^26,27^, and discrete ICNS ganglia are already visible by E16.5 (Extended Data Fig. 3a-d). In contrast, ENS migration starts much earlier at E9.5 from the foregut before the formation of the vagus nerve^28–30^. At E12.5, ENS cells have already colonized most of the GI tract proximal to the cecum and at E14.5 the entire gut; however, only scattered neurons but no visible ganglia are observed at E16.5^14^ (Extended Data Fig. 3e). Our results thus demonstrate that ICNS formation follows a different organizational mechanism from the ENS.

**Fig. 1:**
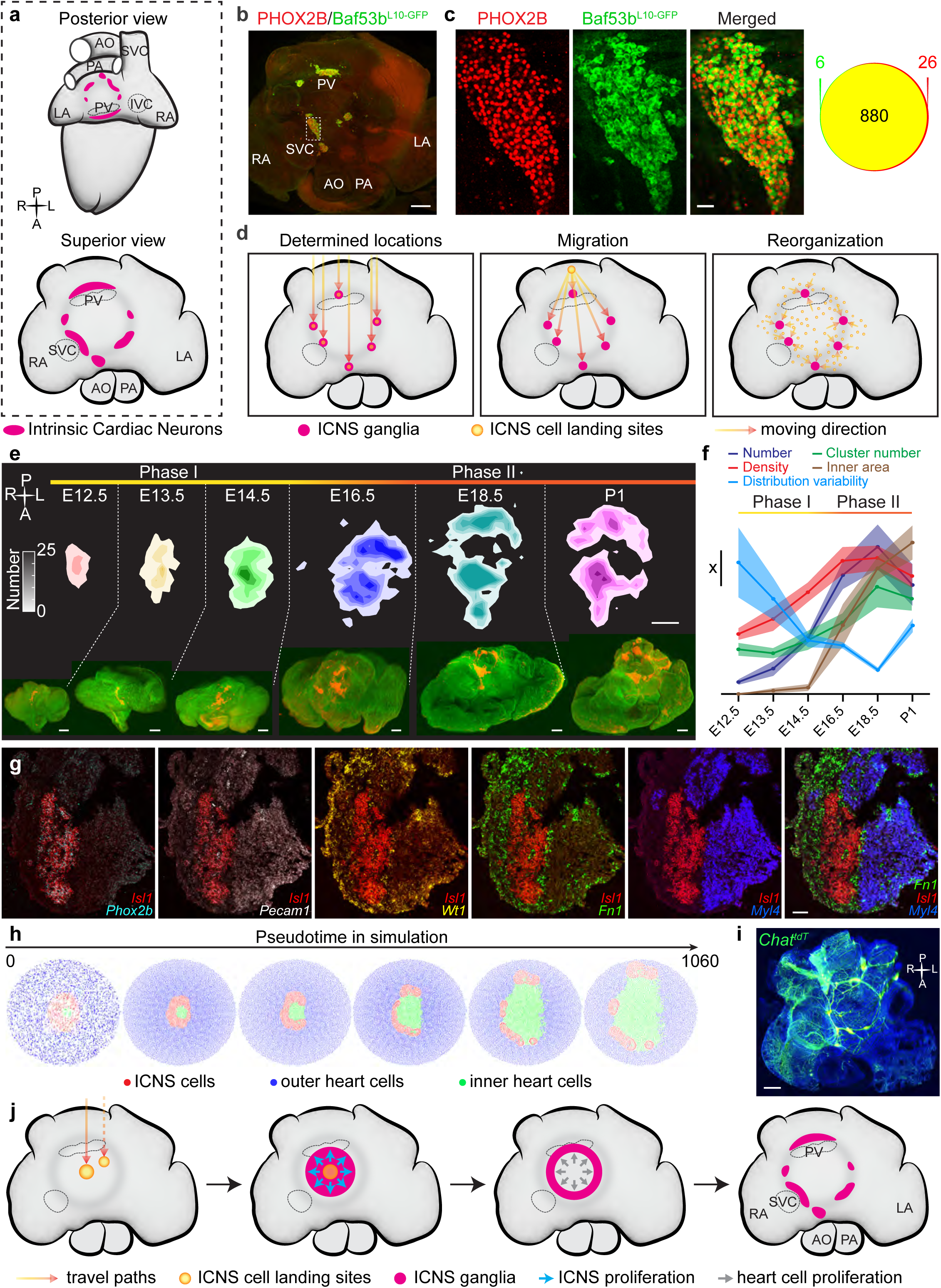
A ‘beads-on-the-necklace’ architecture of the ICNS generated from cardiac cell growth. **a**, Schematic illustration of the ICNS on the mouse heart. LA, left atrium; RA, right atrium; PV, pulmonary veins; IVC, inferior vena cava; PA, pulmonary artery; AO, aorta; SVC, superior vena cava. Magenta dots represent ICNS ganglia. **b**, Superior view of a whole-mount P10 heart from Baf53b^L10-GFP^ mice showing the ICNS labelled with GFP (green) and PHOX2B (red). **c**, Zoom-in images from the dashed region in (**b**). The numbers of single- (red or green) or double-labelled (yellow) cells were counted. **d**, Three models for ICNS formation. **e**, Spatial distributions of PHOX2B^+^ ICNS cells on the heart during development (top). Average numbers across animals are shown in color gradient. Representative images of hearts (superior view) with PHOX2B labelled ICNS cells (red) are shown at bottom. **f**, Quantified ICNS parameters at indicated developmental ages. mean ± sem. x, 500 (number), 0.002 per square micron (density), 1 (cluster number), 30,000 square micron (inner area), 0.014 (distribution variability). **g**, RNAscope Hiplex Assays for indicated genes, showing the ICNS (*Phox2b*^+^/*Isl1*^+^) is physically locked between *Myl4*^+^ cardiomyocytes and *Fn1*^+^ fibroblasts. **h**, simulation of the change of ICNS organization by heart cell proliferation. Red, ICNS cells; green, heart cells in the ICNS inner area; blue, heart cells outside the ICNS ring. **i**, A representative image (superior view) of an adult Chat^tdT^ mouse heart (green), showing the ‘beads-on-the-necklace’ organization of the ICNS. **j**, A model illustrating ICNS formation. Scale bars: 500 μm (**b**), 50 μm (**c, g**), 200 μm (**e**), 1 mm (**i**).

We reason that discrete ICN ganglia could be formed through one of the following hypothesized mechanisms (Fig. 1d): (1) ICNS cells have multiple landing sites, each of which later expands into a ganglion; (2) ICNS cells have one landing site and then actively migrate to their final destinations; or (3) similar as the ENS^14^, ICNS cells are randomly scattered and progressively resolve into individual ganglia. To examine these possibilities, we performed 3D imaging of isolated mouse hearts between E12.5 and postnatal day (P)1 and developed a computational geometric transformation method to quantify the spatiotemporal distribution of ICNS cells across all samples (Fig. 1e-f, Extended Data Fig. 4, see Methods). Because the ICNS is located superficially on the dome of the atrium, this approach allows us to project all ICNS cells onto a 2D plane while preserving their distribution with a negligible z-axis. Our data reveals two anatomical phases of ICNS formation: Phase I (E12.5 – E16.5) which correlates with ICNS landing and proliferation; Phase II (E16.5 - postnatal) that corresponds to the formation of discrete ICNS ganglia. Surprisingly the landing processes are variable among samples: ICNS cells may land on one or multiple (1.25 ± 0.18, n = 12) locations around the center of the atrium (Extended Data Fig. 4e). This variability quickly reduces as loosely connected ICNS cells actively proliferate into a single large ganglion around E13.5-E14.5, with a gradual increase in density and area yet largely unchanged organization. Thus, the initial ICNS landing sites are not correlated with later ICNS ganglia locations. In Phase II, the single ICNS ganglion then undergoes a circular expansion, forming a ring structure that later splits into posterior and anterior groups around E16.5, that eventually localize to a few individual ganglia at P1 which are preserved till adult stage. The number of ICNS ganglia increases as the inner area of the ICNS ring expands while INCDC number, density, and area stay relatively stable until adulthood (Fig. 1e-f, Extended Data Fig. 4f-l).

To determine whether such circular expansion is due to active ICNS migration, we next performed RNAscope-based spatial transcriptomics to examine the cytoarchitecture around the ICNS (Fig. 1g, Extended Data Fig. 5a). Intriguingly, ICNS cells are sandwiched between *Myl4*^+^ cardiomyocytes and *Fn1*^+^/*Wt1*^-^ fibroblasts cells throughout phase II, indicating that they are physically locked in position with minimal migration. Because a migratory wavefront of undifferentiated SOX10^+^ ENS cells is critical for gut colonization^29^ (Extended Data Fig. 5b), we further examined whether a similar structure exists in the developing ICNS. Unlike in the ENS, SOX10^+^/PHOX2B^+^ ICNS cells are not enriched at the edge of the single ICNS ganglion, and their number is extremely low in phase II (Extended Data Fig. 5c), suggesting that a migratory wavefront is absent during ICNS formation and thus not responsible for circular expansion.

We then wonder whether ICNS circular expansion is a passive process determined by mechanical forces from surrounding cardiac cell growth. Indeed, we observed extensive proliferation both inside and outside of the ICNS ring (Extended Data Fig. 5d). We further developed an algorithm to simulate ICNS development under conditions where both inner and outer cardiac cells are rapidly proliferating (see Methods section “**Simulation of ICNS organization”**). Strikingly, this simulation almost completely recapitulates the formation of discrete ICNS ganglia *in vivo* (Fig. 1h, Supplementary Video 1). We also reason that if ICNS circular expansion is caused by rapid cardiac cell growth, the inner area may be wrapped around with extensive nerve fibers but not well innervated. Indeed, we observed a remarkable ‘beads-on-the-necklace’ architecture in Chat^tdT^ mice with almost no tdTomato^+^ innervation to the inner area (Fig. 1i).

Together, our data show that the ICNS and ENS utilize different organization strategies. While ENS colonization relies on migration, ICNS cells are largely stationary after landing. The ‘beads-on-the-necklace’ ICNS architecture is instead achieved through passive circular expansion and tearing from surrounding cardiac cell growth (Fig. 1j). Thus, our study demonstrates the importance of the organ environment in shaping the anatomical organization of organ-intrinsic neurons.

### An intermediate neuronal progenitor state reduces ICNS migration

To further probe the molecular mechanism underlying organ-intrinsic neuron specification, we next sought to uncover the fate landscape of the ICNS through reconstructing the transcriptional trajectory during ICNS development. We first focused on Phase I (proliferation and differentiation) between E12.5 and E16.5 using Wnt1^tdT^ mice in which almost all ICNS cells are fluorescently labelled (Extended Data Fig. 6a). Single-cell RNA sequencing of acutely isolated and fluorescence activated cell sorting (FACS)-purified ICNS cells at various time points (E12.5, E14.5, and E16.5) together reveals a comprehensive genetic landscape of the developing ICNS (Fig. 2a, Extended Data Fig. 6b-c). In total 10,970 *Phox2b*^+^ ICNS cells are profiled using 10x Genomics platform, analyzed using Seurat^31^, and plotted using Uniform Manifold Approximation and Projection (UMAP)^32^ (Fig. 2b-c). Intriguingly, in addition to the classical neuronal developmental states including (1) undifferentiated precursor cells (ICNS-precursors, *Sox10*^+^/*Ednrb^+^*), (2) neuroblast cells (ICNS-neuroblasts, *Ascl1*^high^/*Actl6b*^low^), and (3) maturing neurons (ICNS-neurons, *Sncg*^+^/*Actl6b*^high^), we identified a unique group of dedicated neuronal progenitor cells (ICNS-NPCs, *Sox10*^-^/*Ascl1*^high^/*Mki67*^+^) that is not observed in embryonic ENS cells at similar (Fig. 2d-h, Extended Data Fig. 6d-f, 3,744 ENS cells isolated and sequenced from the small intestine of E14.5 Phox2b^tdT^ mice using 10x Genomics) or earlier/older ages^33,34^. Transcriptional trajectories inferred using RNA velocity^35^ show that in addition to the well-described neural crest-to-neuroblast-to-neuron path as in the ENS^33^, there is an additional fate transition from ICNS-neuroblasts to the intermediate ICNS-NPCs and then to ICNS-neurons (Fig. 2i). Cell-cycle analysis confirms that both ICNS-precursors and ICNS-NPCs are within active cell replication cycles (S/G2/M), while ICNS-neuroblasts and ICNS-neurons are transitioned into the G1 or the post-mitotic G0 phase (Fig. 2j). Immunocytochemistry also demonstrates that the intermediate NPC state is specific to the ICNS as 93.8 ± 3.9 % of proliferating Ki67^+^/PHOX2B^+^ ICNS cells are SOX10^-^ (n = 4) while SOX10^-^/PHOX2B^+^/Ki67^+^ cells are rarely seen in the ENS (10.7 ± 2.0 %, n = 4, Fig. 2k-l). These results reveal that the majority of ICNS-neurons are directly differentiated from SOX10^-^ ICNS-NPCs but not undifferentiated SOX10^+^ ICNS-precursors. Our data thus show that unlike the ENS, the ICNS proliferates using an intermediate progenitor state dedicated for neurons.

**Fig. 2:**
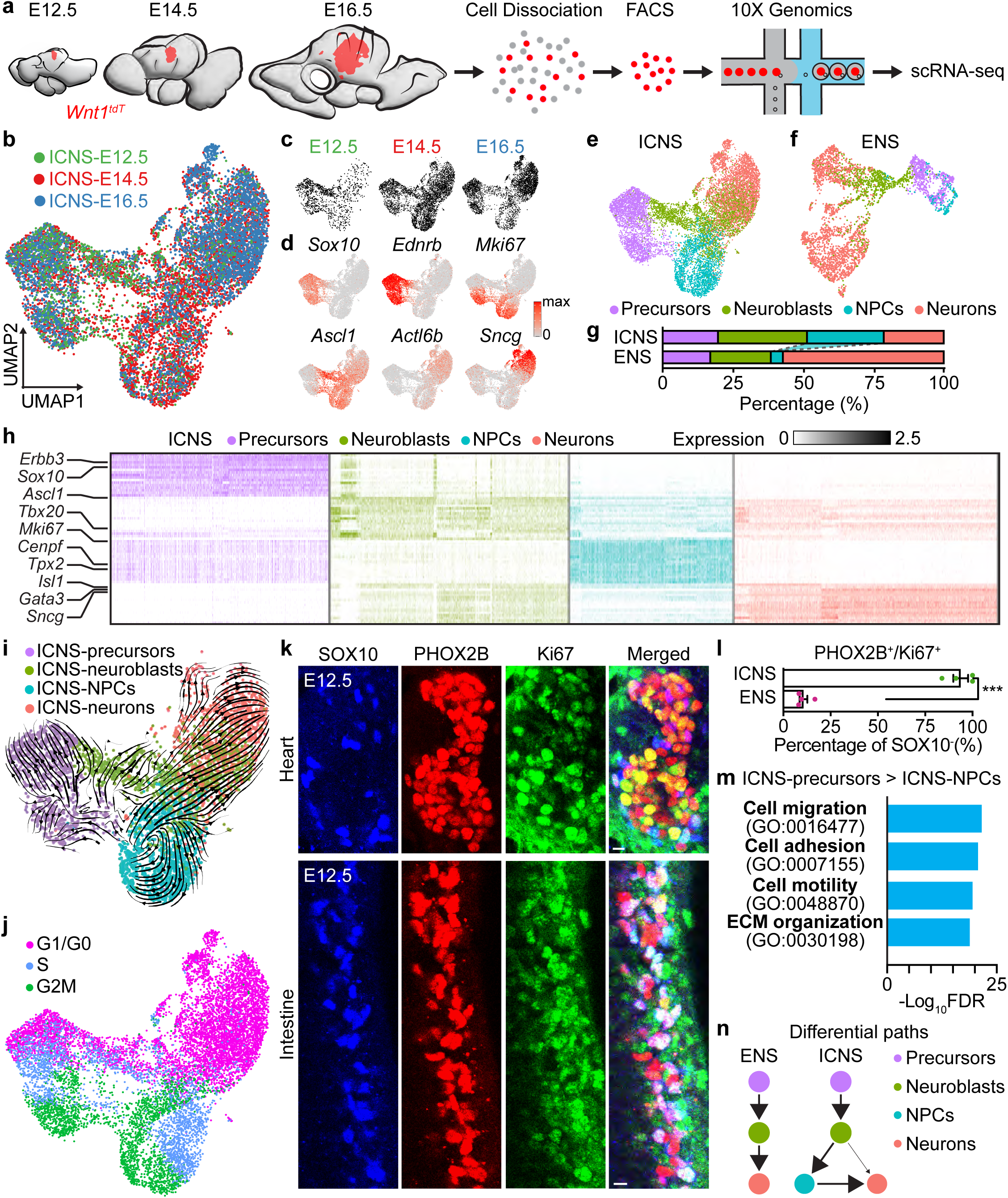
An intermediate neuronal progenitor state reduces ICNS migration. **a**, Schematic illustration of single-cell RNA-sequencing of fluorescently labelled ICNS cells acutely isolated and purified from Wnt1^tdT^ mouse hearts. **b**, UMAP plot of 10970 *Phox2b*^+^ ICNS cells from three developmental ages (color coded). **c**, UMAP plot of ICNS cells from indicated age as in (**b**). **d**, UMAP plot of indicated marker genes for different cell states. **e-f**, UMAP plots of E12.5-E16.5 ICNS cells (**e**) and E14.5 ENS cells (**f**) showing genetically defined cell states (color coded). **g**, Percentage of ICNS and ENS cells, both E14.5, in indicated cell states. **h**, Heat map of genes differentially expressed in various ICNS states (color coded). **i**, UMAP plot of E14.5 ICNS cells showing RNA velocity results (arrows). Cell states are color coded. **j**, UMAP plot of E12.5 – E16.5 ICNS cells, showing cell cycles (color coded). **k**, Representative images of ICNS (top) and ENS (bottom) ganglia labelled with indicated antibodies. Scale bars: 10 μm. **l**, Percentage of SOX negative proliferating precursors (PHOX2B^+^/Ki67^+^) in ICNS and ENS. **m**, Selected top GO pathways of DEGs in ICNS-precursors over ICNS-NPCs. FDR, false discovery rate. **n**, Models illustrating differential paths for ENS and ICNS generation.

We next compared differentially expressed genes (DEGs) among ICNS states to understand the biological differences during state transitions. Interestingly, we discovered that cell migration, cell motility, and extracellular matrix (ECM) organization are the top Gene Ontology (GO)^36,37^ biological processes enriched in ICNS-precursors that are downregulated in ICNS-NPCs and other ICNS states (Fig. 2m, Extended Data Fig. 7). We then performed trajectory analyses using Monocle 3^38^ to examine gene dynamics during state transitions across the neurogenesis phase (Extended Data Fig. 8). Using hierarchical clustering^39^, we grouped highly variable genes along the target trajectory into distinct sets (modules) based on their pseudo-temporal expression patterns. The physiological function of each gene module is inferred with GO analysis. Our data show that the ICNS-precursor to ICNS-neuroblast transition begins with downregulation of genes featuring neural crest-derived precursor cells^40^ and coincidentally upregulation of genes involved in RNA processing and biosynthesis (Extended Data Fig. 8). This is followed by upregulation of genes involved in neurogenesis and finally genes for synaptic transmission and neuron projection development. Our analysis thus reveals four key transition steps between ICNS-precursors and ICNS-neuroblasts. Interestingly, our study demonstrates that cell motility is immediately downregulated when ICNS-precursors differentiate (Extended Data Fig. 8c), including many genes critically involved in neural crest cell migration and ENS colonization^29,41–43^ such as *Sox10*, *Ednrb*, *Itgb1*, *Tnc*, and *Dusp6*. Cell motility score calculated using the corresponding GO term (GO:0048870) also dramatically decreases in all other ICNS cell states compared to undifferentiated ICNS-precursors (Extended Data Fig. 8d), suggesting an instant loss of migratory capability of ICNS cells after landing on the heart. These findings together provide a molecular explanation for our anatomical observation that while SOX10^+^ ENS cells are highly migratory at the wavefront and are essential for gut colonization^29^, there is minimum migration during ICNS formation (Fig. 1). Our results thus show that the ICNS employs an intermediate NPC state for effective proliferation without migration to better accommodate its organ environment (Fig. 2n).

### Extracellular matrix anchors ICNS precursors at the correct heart location

We reason that before differentiation, ICNS-precursors should still have the ability of migrating along the vagus nerve to more distal heart locations such as the ventricles, similar to the ENS cells colonizing the entire gut. However, we observe that they always settle around the center of the atrium, suggesting that there must be attractive signals from the future ICNS area. To determine the molecular mechanisms that immobilize ICNS-precursors, we isolated cardiac cells surrounding (heart^near^) or further away (heart^far^) from the ICNS from E14.5 mouse embryos and performed single-cell RNA-sequencing (Fig. 3a-b). In total 13,904 cells (8,260 from heart^near^ and 5,644 from heart^far^) were sequenced using the 10x Genomics platform, integrated using Seurat, and clustered using classical cardiac gene markers^44,45^ into cardiomyocytes (CM, *Tnni3*), fibroblasts (FB, *Dcn*), endothelial cells (EC, *Pecam1*), epicardial cells (EP, *Wt1*), myeloid cells (MY, *C1qa*), and ICNS cells (ICNS, *Phox2b*). As expected, only heart^near^ but not heart^far^ contains ICNS cells (Fig. 3c-d). Two major cardiac cell types (cardiomyocytes and fibroblasts) show significant transcriptional shifts with location-specific gene signatures (Fig. 3b-c). Moreover, the percentage of fibroblasts almost doubles in heart^near^ over heart^far^ (Fig. 3d), indicating that they are highly enriched around the ICNS area. Cell-cell communication analysis using CellChat^46^ also predicts that fibroblasts and cardiomyocytes are the top 2 sources for signalling to ICNS (Fig. 3e). Spatially, cardiomyocytes and fibroblasts immediately contact and wrap around ICNS cells during development (Fig. 1g). These results together suggest that fibroblasts and cardiomyocytes may provide critical cell-cell interaction signals to attract and immobilize ICNS cells.

**Fig. 3:**
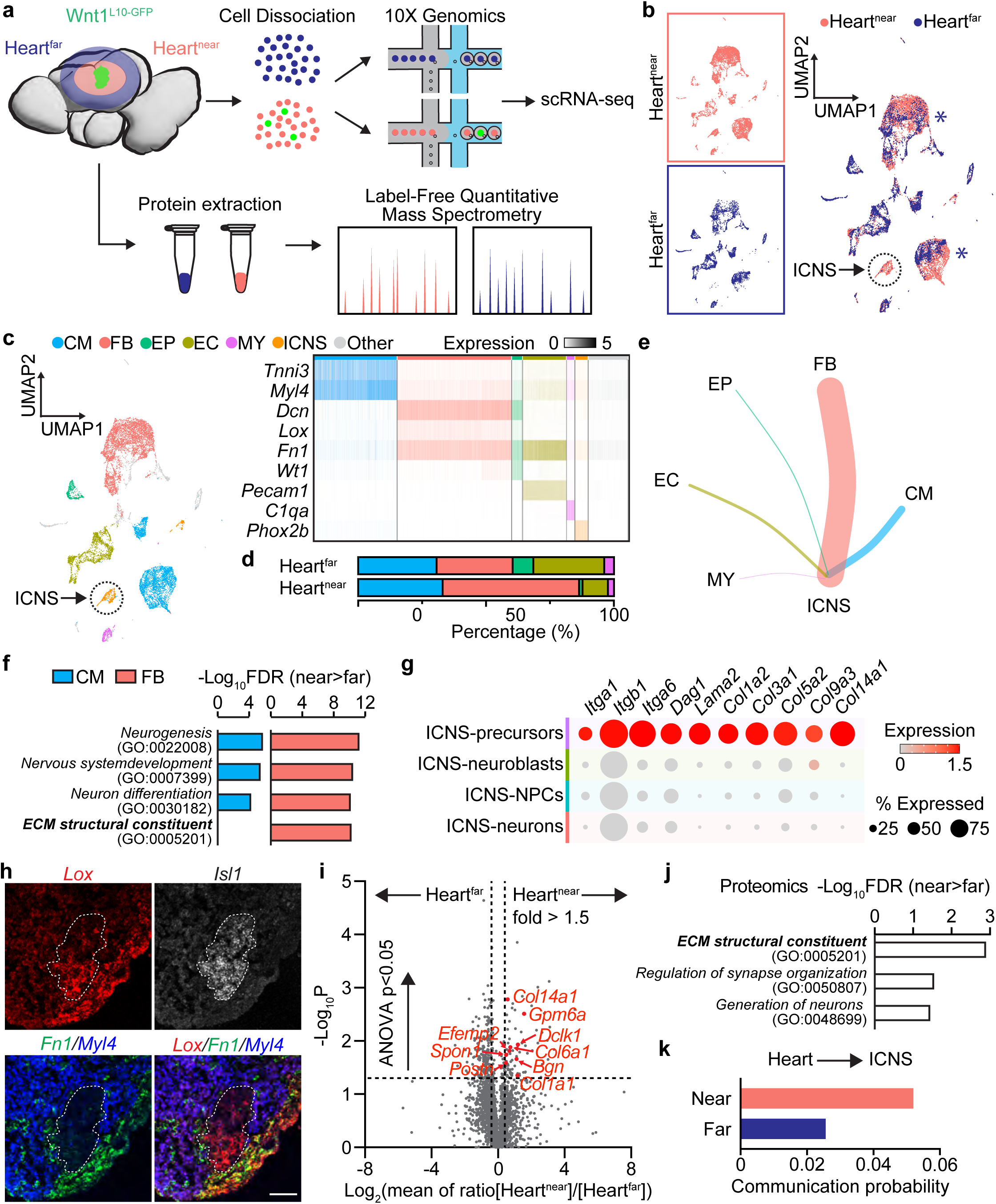
Extracellular Matrix (ECM) anchors ICNS-precursors at the correct heart location. **a**, Schematic illustration of single-cell RNA-sequencing and proteomic analyses of heart regions surrounding (heart^near^) or away (heart^far^) from the ICNS at E14.5. **b**, UMAP plots of E14.5 heart cells integrated from both heart^near^ and heart^far^ (color coded). The dashed circle indicates the ICNS. Asterisks indicate transcriptionally shifted cell types. **c**, UMAP plot of heart cells as in (**b**), colored by genetically defined cell types (left) and the heatmap of cell-type specific marker genes (right). CM, cardiomyocytes; FB, fibroblasts; EP, epicardial cells; EC, endothelial cells; MY, myloid cells; ICNS, intrinsic cardiac nervous system. **d**, Percentages of different heart cell types in heart^near^ and heart^far^ datasets. **e**, Communication probabilities from CellChat analysis (shown as line thickness) between various E14.5 heart cell types as donors and ICNS cells as the recipient. **f**, Selected top GO pathways of cardiomyocyte (CM, blue) or fibroblast (FB, coral) DEGs in heart^near^ over heart^far^. FDR, false discovery rate. **g**, Dot plot of indicated ECM genes between various ICNS cell states (color coded). **h**, RNAscope HiPlex analysis of indicated genes showing that *Lox* is jointly expressed between the ICNS (*Isl1*^+^, dashed circle) and fibroblasts (*Fn1*^+^). Scale bar: 50 μm. **i**, Volcano plot of differences in abundance in the proteome of heart^near^ and heart^far^. Dashed horizontal line, ANOVA p = 0.05. Dashed vertical lines, fold change = ± 1.5. Selected ECM proteins enriched in heart^near^ are highlighted in red (showing encoding genes). **j**, Selected top GO pathways of proteins enriched in heart^near^ over heart^far^. FDR, false discovery rate. **k**, Communication probabilities from CellChat analysis between different heart^near^ and heart^far^ cell types as donors and ICNS cells the recipient.

We therefore focused on cardiomyocytes and fibroblasts and analyzed their DEGs between heart^near^ and heart^far^. Strikingly, in both cardiomyocytes and fibroblasts, heart^near^ enriched genes are highly associated with neurogenesis, nervous system development, and neuron differentiation (Fig. 3f). These include genes encoding both secreted signalling molecules such as *Sema3d*, *Inhba*, and *Igf1* and cell-cell contact signals such as *Dlk1*, *Thbs2*, *Nrxn3*, *Ncam1*, and *Sema5a/b* (Extended Data Fig. 9a). Furthermore, one top GO molecular function of heart^near^ fibroblast DEGs is ECM structural constituent (GO:0005201), and compared to heart^far^, genes encoding specific laminin, elastin, and collagen isoforms including *Lama2*, *Eln*, *Col1a1*, *Col1a2*, *Col6a1*, and *Col14a1* are uniquely expressed or upregulated in heart^near^ fibroblasts (Fig. 3f, Extended Data Fig. 9a). Intriguingly, our sequencing data also discovers that ECM genes are highly expressed in ICNS cells, in particular in ICNS-precursors (Fig. 3g, Extended Data Fig. 9b). Moreover, RNAscope *in situ* hybridization analysis reveals that *Lox*, a copper-dependent amine oxidase that covalently crosslink and stabilize ECM molecules^47^, is jointly expressed at the fibroblast-ICNS border (Fig. 3h), suggesting that in addition to a neuro-attractive role, cardiac fibroblasts may also help secure ICNS cells in position through physical ECM interactions.

To better visualize the spatial expression of heart^near^ DEGs, we next performed 10x Visium based spatial transcriptomics of E14.5 hearts. Because 10x Visium does not provide single-cell resolution, we used Wnt1^Sun1-GFP^ mice to facilitate the identification of ICNS locations. Consistent with single-cell RNA-sequencing results, our spatial transcriptomics data show that neuronal attractive cues and ECMs are preferentially expressed around the ICNS area (Extended Data Fig. 10). To examine at the protein level, we further performed Label-Free Quantitative Mass Spectrometry of heart^near^ and heart^far^ samples isolated from E14.5 Wnt1^L10-GFP^ embryos (Fig. 3a). In total 108 heart^near^ enriched proteins are identified (ANOVA p < 0.05), most of which are neuronal markers highly associated with synaptic structures, neurofilament organization, neuron morphogenesis, and other neuronal functions, demonstrating the validity of the result (Fig. 3i-j, Extended Data Fig. 11). Interestingly, our proteomic data again highlights ECM structural constituent as one of the top GO molecular functions of heart^near^ enriched proteins, including collagens such as CO1A1, CO6A1, and CO14A1 and cell adhesion molecules such as POSTN and BGN (Fig. 3i-j). Our study therefore demonstrates that cell-cell contact signals and ECMs are preferentially expressed around the ICNS area at both transcriptomic and protein levels.

We then performed CellChat analysis to characterize cell-cell communications between heart and ICNS cells. Our result shows that indeed the calculated communication probability (which infers the interaction strength) of heart^near^-ICNS (donor-recipient) is higher than heart^far^-ICNS (Fig. 3k). Moreover, CellChat analysis further underlines THBS, NCAM, IGF, SEMA3, SEMA5, ACTIVIN, LAMININ and COLLAGEN pathways in heart-ICNS communications (Extended Data Fig. 12). Interestingly, ECM-interactions including LAMININ and COLLAGEN pathways show highest probability at ICNS-precursor states, between cardiac fibroblast and ICNS-precursors, consistent with the idea that ECM interactions help anchor ICNS-precursors at the appropriate heart location. On the other hand, communication probabilities of THBS, NCAM, IGF, SEMA3, SEMA5, ACTIVIN and NRXN pathways increase as ICNS-precursors differentiate into ICNS-neurons, suggesting that organ cues are also crucial for neuron maturation. In summary, our spatial studies highlight the role of ECMs in ICNS formation.

### ICNS and ENS have similar precursors but diverse differentiation paths

We next seek to understand whether organ-intrinsic neurons have common or unique differentiation paths to become dedicated for their target organs. To address this question, we performed single-cell RNA sequencing of maturing ICNS-neurons acutely isolated from Baf53b^L10-GFP^ mice at E16.5, E18.5, and P1 to better understand ICNS maturation (Phase II, Extended Data Fig. 13a-d). Our analysis reveals three major ICNS-neuron subtypes differentially labelled by *Ddah1*^+^, *Npy*^+^, and *Vip*^+^ (Extended Data Fig. 13e-f), suggesting that the ICNS is less heterogeneous than the ENS. We further performed RNAscope HiPlex Assay to examine the spatial distribution of ICNS-neuron subtypes (Extended Data Fig. 13g). Our data shows that consistent with single-cell RNA-sequencing data, *Ddah1*, *Npy*, and *Vip* are expressed in largely separate but intermingled ICNS-neurons at E16.5, reminiscent of the salt-and-pepper distribution pattern of neuron subtypes within the ENS.

To reveal ENS differentiation at a high temporal resolution, we leveraged existing high quality ENS datasets^33,34^ at E9.5, E10.5, E11.5, E15.5, E18.5 and performed additional single-cell RNA sequencing of ENS cells at E13.5 and E14.5 using Phox2b^tdT^ mice (Extended Data Fig. 14a-e). We then integrated ICNS and ENS single-cell datasets at various embryonic stages (10,441 ICNS cells from E12.5 (1,124), E14.5 (3,744), E16.5 (4,615), E18.5 (958); 14,979 ENS cells from E9.5 (191), E10.5 (369), E11.5 (463), E13.5 (3,500), E14.5 (4,395), E15.5 (3,354), E18.5 (2,707)) to compare differentiation paths between the ICNS and ENS (Fig. 4a, Extended Data Fig. 14f). Consistent with the view that organ-intrinsic neurons are specialized for their target organs, mature ICNS- and ENS-neurons express different sets of transcription factors, surface receptors, and neurochemicals (Fig. 4b). For example, *Isl1* and *Hoxa5* differentially mark ICNS- and ENS-neurons in RNAscope HiPlex Assay (Fig. 4c). As shown earlier (Fig. 2f), the NPC population prominent in the ICNS is almost absent in the ENS (Fig. 4d). Otherwise, ICNS- and ENS-precursor and neuroblast cells show almost perfect overlap on the UMAP, demonstrating that they are highly similar in genetics. Yet their fates start to separate immediately as they transition into maturing neurons. Interestingly, while the binary fate split of ENS-neurons happens right after neurogenesis, the ICNS-neuron differentiation trajectory sits right in between the two ENS-neuron branches and its fate selection begins at a much later pseudo-time (Fig. 4e), suggesting that the differential potential of ICNS cells is limited compared to the ENS cells. This result is consistent with and may partially explain our notion that mature ICNS-neurons are much less heterogeneous than ENS-neurons (Extended Data Fig. 13).

**Fig. 4:**
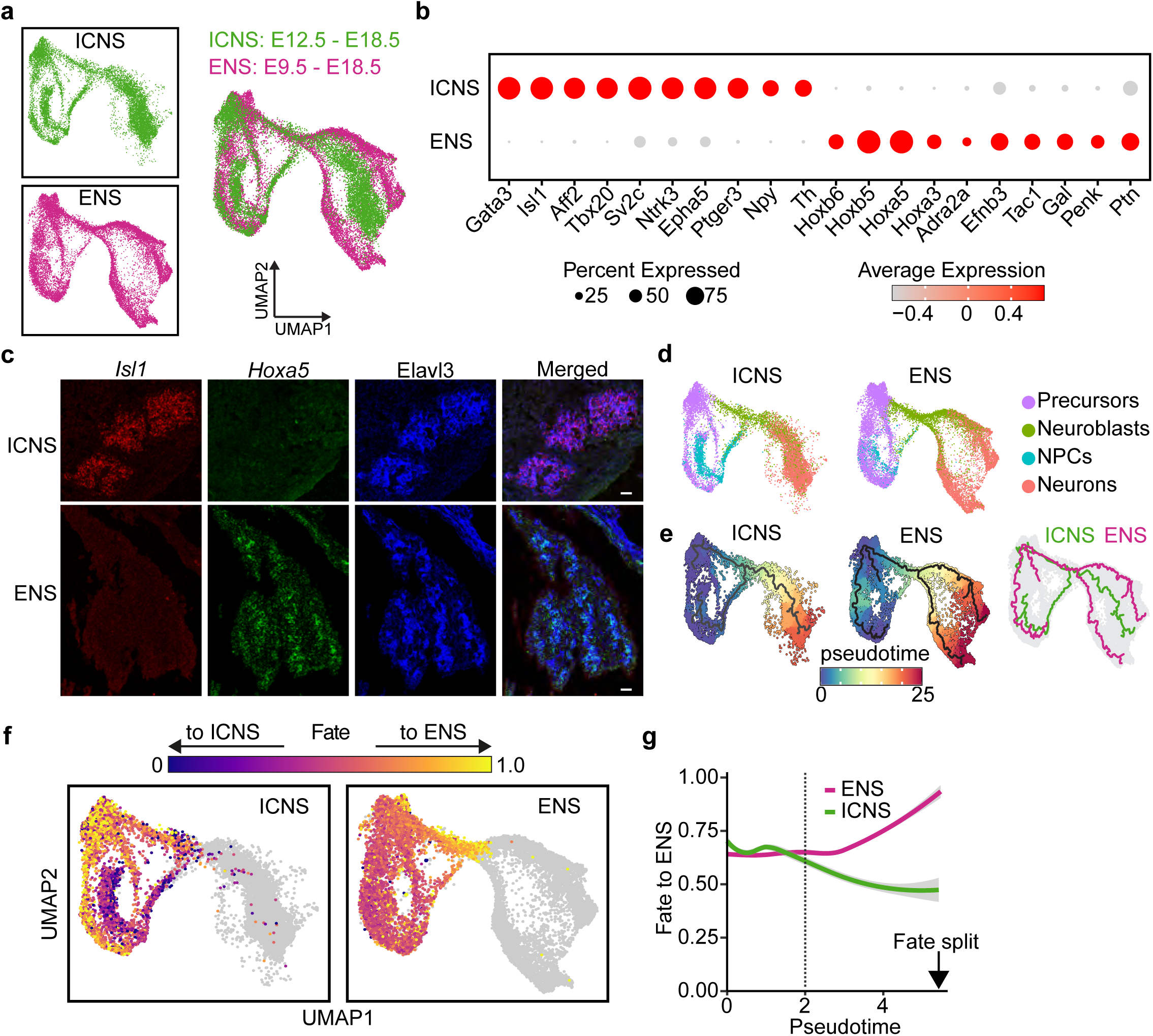
ICNS and ENS have similar precursors but diverse differentiation paths. **a**, UMAP plot of integrated ICNS (10,441 cells from E12.5, E14.5, E16.5, and E18.5, green) and ENS (14,979 cells from E9.5, E10.5, E11.5, E13.5, E14.5, E15.5, and E18.5, magenta) cells. **b**, Dot plots of transcription factors, surface receptors, and neurochemicals differentially expressed in ICNS and ENS cells. **c**, RNAscope HiPlex analysis of *Isl1* (red) and *Hoxa5* (green) in ICNS and ENS cells. All neurons are revealed using Elavl3 staining (blue). **d-f**, UMAP plots of ICNS and ENS cells as in (**a**), showing (**d**) different cell states (color coded), (**e**) differentiation trajectories (left, black trace; right, green for ICNS, magenta for ENS) and pseudotime (left, colored) predicted using Monocle 3, and (**f**) fate probability (0 - 1) to become ENS-neurons (color coded). **g**, Fate probability to ENS-neurons of ICNS- (green) and ENS- (magenta) precursors/neuroblasts along the predicted pesudotime. Fitted by loess (locally estimated scatterplot smoothing), mean ± 95 % ci. The dashed line marks the boundary between precursor and neuroblast cells.

To better understand organ-intrinsic neuron fate selection, we further performed scTIE, a method that infers regulatory relationships predictive of cellular state changes from large single-cell temporal datasets^48^. Interestingly, scTIE results show that both ENS- and ICNS-precursors are possible of becoming ENS- or ICNS-neurons, and their fate probabilities gradually shift to their corresponding neuron types during neurogenesis (Fig. 4f-g). As ICNS-neuroblasts transition to the unique ICNS-NPCs, their fate already switches to ICNS-neurons over ENS-neurons. Not surprisingly, scTIE analysis highlights the role of transcription factors in organ-intrinsic neuron fate selection. Indeed, the expression of many transcription factors specific for ICNS-neurons (such as *Gata3*, *Isl1,* and *Tbx20*) and ENS-neurons (such as *Hoxa5* and *Hoxb5*) starts to rise during the precursor-to-neuroblast transition (Extended Data Fig. 14g-h). Together, through cross-organ comparisons, our results demonstrate that visceral organs regulate the differentiation path of organ-intrinsic neurons and control their fate selection.

### Visceral organs influence the identity of organ-intrinsic neurons through ECM-receptor interactions

To better understand how visceral organs impact organ-intrinsic neuron fate selection, we next sought to characterize in more detail cell-cell interactions between visceral organs and their intrinsic neurons, in particular precursors and neuroblasts. To achieve this, we further performed single-cell RNA sequencing of acutely isolated intestine cells from E14.5 mice (Extended Data Fig. 15a) and merged this new dataset with our existing single-cell dataset of E14.5 heart^near^ cells, and integrated ICNS and ENS datasets. We then performed CellChat analysis to compare ligand-receptor interactions between heart^near^-ICNS and gut-ENS pairs (Fig. 5a). Interestingly, among the three categories of molecular interactions (cell-cell contact, secreted signalling, and ECM-receptor), heart-ICNS and gut-ENS have similar communication probabilities for cell-cell contact and secreted signalling interactions, involving predominantly the same ligand-receptor pairs (Fig. 5b). Many of these interactions are general pathways associated with neuronal maturation such as SEMA3, NCAM, and IGF (Extended Data Fig. 15b). In contrast, heart-ICNS has a much higher (> 4 fold) communication probability for ECM-receptor interaction than gut-ENS (Fig. 5b). Some ligand-receptor pairs are shared between heart-ICNS and gut-ENS but are much stronger in heart-ICNS. Others are unique for heart-ICNS. Detailed analysis highlights a number of ECM-receptor pathways in heart-ICNS that are with much lower communication probability or almost absent in gut-ENS, including THBS, COLLAGEN, and LAMININ (Fig. 5c). Our results thus demonstrates that ECM-receptor interaction represents the major difference between heart-ICNS and gut-ENS communications. To further understand whether such difference in the interaction between visceral organs and their intrinsic neurons is from visceral organ cells (donor), organ-intrinsic neurons (recipient), or both, we swapped the donors and recipients and re-performed the analysis for heart-ENS and gut-ICNS (Fig. 5a). Consistent with the notion that most of the cell-cell contact and secreted signalling interactions are shared between heart-ICNS and gut-ENS, swapping donors and recipients (meaning heart-ENS and gut-ICNS) does not change the communication probabilities of these two categories (Fig. 5d). Surprisingly, for ECM-receptor interaction, both heart-ENS and gut-ICNS have higher communication probabilities than gut-ENS (Fig. 5d, 1.8- and 2.8-fold respectively). Both ICNS and ENS have higher communication probabilities with heart than gut (1.6- and 1.8-fold receptively). Meanwhile, both heart and gut have higher communication probabilities with ICNS than ENS (2.4- and 2.8-fold respectively). These results suggest that both donors and recipients contribute to the ECM-receptor interaction difference between heart-ICNS and gut-ENS, and the impact from recipients (organ-intrinsic neurons) is more prominent. Indeed, many ECM genes are differentially expressed in ICNS cells over ENS cells, in particular in ICNS-precursors (Extended Data Fig. 15c). Our data together indicate that visceral organs likely regulate organ-intrinsic neurons’ fate selection through ECM-receptor interactions.

**Fig. 5:**
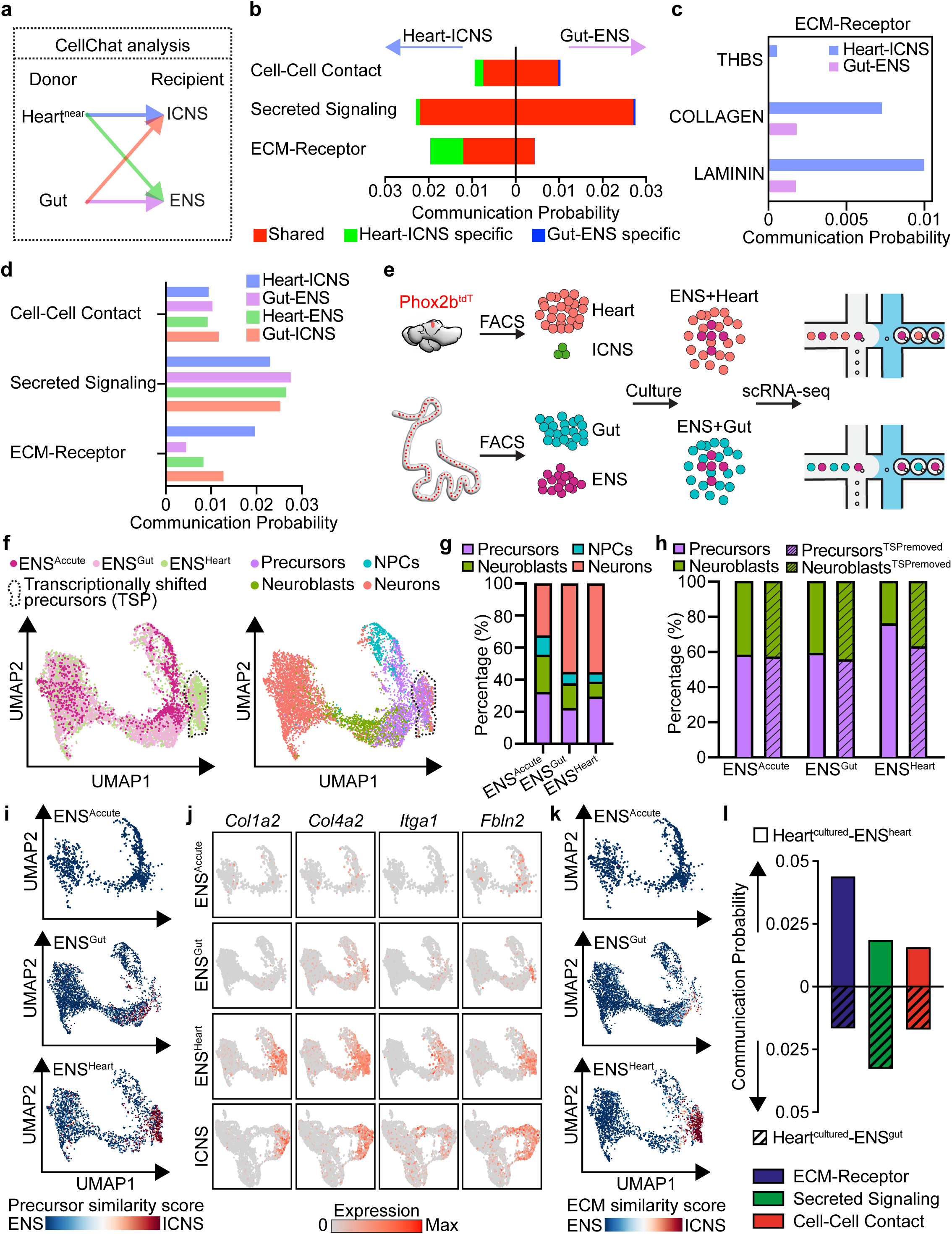
Visceral organs influence the identity of organ-intrinsic neurons through ECM-receptor interactions. **a**, Schematic illustration of CellChat analyses using donor-recipient pairs (heart^near^ or gut cells as donors, ICNS or ENS as recipients). **b**, Communication probabilities of heart-ICNS (left) and gut-ENS (right) donor-recipient pairs, among the three categories of molecular interactions (cell-cell contact, secreted signalling, and ECM-receptor). Cumulative contributions from heart-ICNS specific, gut-ENS specific, and shared ligand-receptor pairs are color coded. **c**, Communication probability of indicated ECM-receptor pathways that show stronger interactions in heart-ICNS over gut-ENS. **d**, Communication probabilities of indicated donor-recipient pairs (color coded), among the three categories of molecular interactions. **e**, Schematic illustration of ENS co-cultures with heart or gut cells. **f**, UMAP plot of ENS cells (ENS^acute^, 1,000, acutely isolated; ENS^gut^, 3,768, co-cultured with gut cells; ENS^heart^, 2,725, co-cultured with heart cells), colored by condition (left) and cell states (right). The dashed circle shows a transcriptionally shifted subset of non-proliferating precursor cells when co-cultured with heart cells. **g-h**, Cumulative percentage of ENS cells (acutely isolated, co-cultured with heart or gut cells) in all four cell states (**g**) and between precursor/neuroblast states (**h**) with (plain) and without (stripes) the transcriptionally shifted subset of non-proliferating precursor cells as shown in (**f**). **i**, UMAP plots of ENS cells at indicated conditions, showing precursor similarity score towards ICNS calculated using ICNS-precursor DEGs (see Methods). **j**, UMAP plots of ENS^acute^, ENS^gut^, ENS^heart^, and ICNS (E14.5) colored by expression of indicated ECM genes. **k**, UMAP plots of ENS cells as in (**i**), showing ECM similarity score towards ICNS calculated using ICNS-precursor ECM genes. l, Communication probabilities of heart^clutured^-ENS^heart^ (top, plane) and heart^clutured^-ENS^gut^ (bottom, strips) donor-recipient pairs, among the three categories of molecular interactions (ECM-receptor, blue; secreted signalling, green, and cell-cell contact, red).

We thus ask whether changing the organ environment would affect the identity of organ-intrinsic neurons. To test this, we isolated fluorescently labelled ENS cells from E13.5 Phox2b^tdT^ mice and co-cultured them with fluorescence-negative organ cells isolated from the gut or heart of the same animals. After 4-5 days, co-cultured cells were dissociated and subjected to single-cell RNA sequencing using 10x Genomics platform (Fig. 5e). Co-cultured ENS cells were identified as *Phox2b*^+^ clusters from unsupervised clustering after quality control (Fig. 5f, 3,768 from ENS co-cultured with gut, ENS^gut^; 2,725 from ENS co-culture with heart, ENS^heart^). Both ENS^gut^ and ENS^heart^ cells have clear precursor-, neuroblast- and maturing neuron-states. Compared to acutely isolated ENS cells at E13.5, ENS^gut^ cells have a higher percentage of maturing neurons but comparable precursor-neuroblast, demonstrating that the co-cultured ENS has a lower proliferation rate *in vitro* but retained the capacity for neuron differentiation (Fig. 5g-h). ENS^heart^ cells have similar percentage of maturing neurons as ENS^gut^ cells, indicating that neuronal maturation from neuroblasts is not affected (Fig. 5g). On the other hand, ENS^heart^ cells have fewer neuroblasts and a much higher precursor-neuroblast ratio, suggesting a reduction in precursor differentiation (Fig. 5h). Intriguingly, we noticed that co-culturing with heart causes transcriptional shift in a subset of non-proliferating precursor cells (Fig. 5f), while the rest of ENS^heart^ precursors and neuroblasts have a comparable ratio with ENS^gut^ and ENS acutely isolated at E13.5 (Fig. 5h). Our data thus demonstrates that co-culturing with heart cells changes genetic signatures of ENS-precursors, raising the possibility that if the identity of these affected cells might shift towards the ICNS. We then calculated an ICNS-ENS precursor similarity score using the top 400 DEGs between ICNS- and ENS-precursor cells (200 for each direction). Our results show that the transcriptionally shifted ENS^heart^ population dramatic transitions to the ICNS-precursor state (Fig. 5i). GO analysis of DEGs between ENS^heart^ and ENS^gut^ non-proliferating precursors reveals, in addition to gene modules involved in various developmental processes, the role of ECM and cell adhesion genes during this transition (Extended Data Fig. 15d). Strikingly, a number of ECM genes preferentially expressed in ICNS-precursors including *Col1a2*, *Col4a2*, *Itga1*, and *Fbln2* are induced in ENS^heart^ cells (Fig. 5j). ICNS-ENS ECM similarity score, calculated using ECM genes differentially expressed between ICNS and ENS precursor cells, further confirms that the transcriptionally shifted ENS^heart^ population selectively expresses ICNS-like ECM genes (Fig. 5k). Moreover, CellChat analysis shows that compared to ENS^gut^ cells, ENS^heart^ cells have stronger communications with co-cultured heart cells, in particular via ECM-receptor interactions (Fig. 5l), reminiscent the differences between heart-ICNS and gut-ENS pairs (Fig. 5b). Together, our results demonstrate that visceral organs can influence the identity and differentiation of organ-intrinsic neurons through regulating their ECM environments.

## Discussion

As a key component of the body-brain axis, the importance of organ-intrinsic neurons in organ pathophysiology, interoception, and cognition has been increasingly appreciated. Appropriate specialization of organ-intrinsic neurons for their host organs in both molecular architecture and anatomical organization is a critical process yet the underlying mechanisms remain largely mysterious. Integrating high-resolution whole-organ imaging and single-cell genomics approaches, we describe how ICNS formation is shaped by the heart via interactions with surrounding cardiac cells. Through comparisons with the ENS on the gut, our cross-organ study further uncovers the differential strategies that visceral organs use to drive the specialization of their organ-intrinsic nervous systems.

Although developed from the same origin^13^, organ-intrinsic neurons need to select their fates and change their organization patterns to adapt to their host organs for optimized performance. For example, the ENS colonizes the entire GI tract to provide sensory and regulatory signals at high spatial resolution^7^. On the other hand, extreme mechanical forces during intense contractions make the heart particularly hostile for neurons. Unlike the ENS, the ICNS dose not colonize the entire heart. Instead, it is located around the apical center of the atria, one of the regions least affected by heart contractions. Our study shows that the heart plays an active role in shaping the organization of the ICNS. First, the heart traps ICNS-precursors at the right spot through an extensive network of ECM-receptor interactions (Fig. 3). Intriguingly, even this step might be triggered by the heart as when co-cultured in vitro, cardiac cells are able to induce the expression of corresponding ECM genes/receptors in ENS-precursors that normally do not express these ECM genes (Fig. 5). Second, the ICNS uses an intermediate dedicated neuronal progenitor state that has almost no migratory capacity instead of the highly migratable ICNS-precursors to produce appropriate number of ICNS-neurons (Fig. 2). This fate convergence^49^ further secures the ICNS at the optimized location. Third, as ICNS cells proliferate into a large ganglion, the rapid growth of surrounding heart cells passively terminates ICNS proliferation, separates the single ganglion apart into several smaller ganglia, and delivers them to separate designated locations (Fig. 1). Together, this interesting strategy effectively ensures a stereotypical ICNS organization without the need for precise regulation of the number of ICNS cells landed on the heart via the vagus nerve, their landing sites, or even their identities.

Organ-intrinsic neurons are closely associated with not only autonomic dysfunctions but also neurological disorders such as Alzheimer’s disease and Parkinson’s disease^50,51^. Looking forward, cross-organ comparisons using high-throughput large-scale systems approaches promise to provide a deeper understanding of the differential developmental blueprints of diverse organ-intrinsic nervous systems and inspire innovative therapies for disease treatment.

## Acknowledgements

We thank N. Palm, E. Gracheva, and P. De Camilli for sharing mice and equipment, Gefei Wang for advice on spatial transcriptomics analysis, S. Wilson, L. Shao, and Yale CNNR Imaging Core for assistance with microscopy, Guilin Wang and YCGA for assistance with single-cell RNA-sequencing and 10x Visium spatial transcriptomics, L. TuKiet and MS & Proteomics Resource at Yale University for providing the necessary mass spectrometers and the accompany biotechnology tools, D. Trotta and the Yale Flow Cytometry Facility for assistance with FACS, J. Greenwood and Yale Neuroscience Technology Core for making imaging apparatus and chambers. Funding was provided by NIH (R01HL150449, DP2HL151354 to R.B.C; R01HL148008 to L.H.Y.), the G. Harold & Leila Y. Mathers Foundation (R.B.C.), the Aligning Science Across Parkinson’s (ASAP) initiative (R.B.C. and L.Z.), the Allen Discovery Center program, a Paul G. Allen Frontiers Group advised program of the Paul G. Allen Family Foundation (R.B.C. and L.Z.), and Kavli Postdoctoral Fellowship (I.H.).

## Author Contribution

I.H., L.Z., and R.B.C. designed experiments, analyzed data, and wrote the manuscript. I.H. led and performed all experiments. Y.L. performed scTIE and RNA velocity analyses. I.H. and Y.G. performed Monocle trajectory analysis. Q.J.X. pioneered ICNS isolation, purification, and sequencing. Y.S. helped ICNS image analysis. I.H., R.W., D.Y., J.Z., and L.Z. performed 10x Visium spatial transcriptomics and analysis. I.H. performed all other data analyses. L.H.Y. and R.B.C. supervised heart anatomical studies. H.Z. and L.Z. supervised bioinformatic studies.

## Data and Code Availability

Single-cell RNA-sequencing and 10x Visium based spatial transcriptomics data generated for the current study will be made publicly available at NCBI Gene Expression Omnibus (GEO). Proteomics data will be made publicly available at the PRIDE repository. Code for Seurat, scTIE, Monocle 3, and CellChat will be available at github.com. Additional data or code related to this study may be available upon request.

## Extended Data Figure Legends

**Extended Data Fig. 1:**
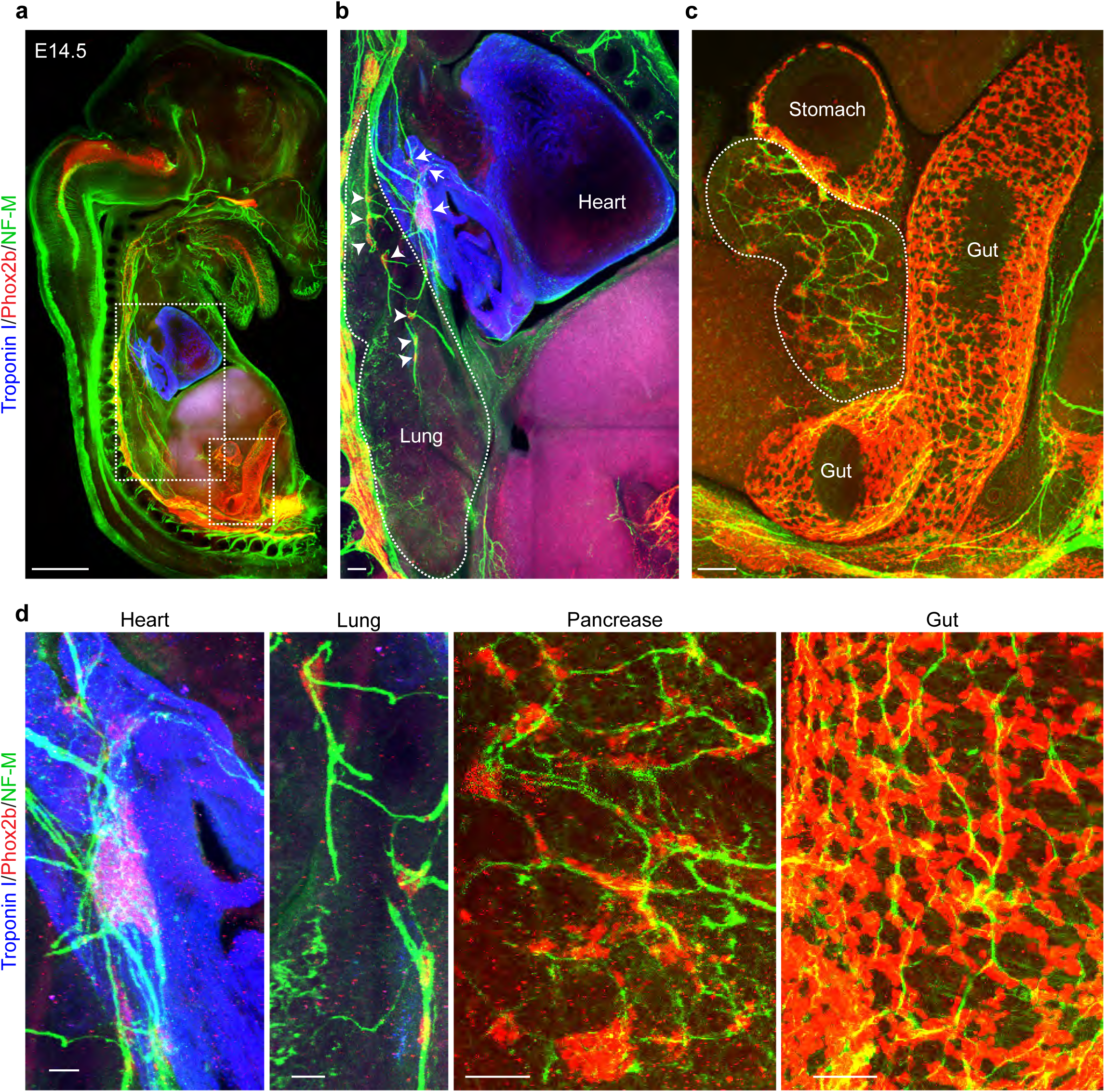
Diverse organ-intrinsic nervous systems. **a**, Whole-embryo 3D imaging of a cleared E14.5 mouse embryo stained with Phox2b (red, organ-intrinsic neurons), Neurofilament M (NF-M, green, nerves), and Troponin I (blue, heart). **b-c**, Zoom-in images from the dashed region in (**a**), showing ICNS cells (**b**, arrows), intrapulmonary neurons (**b**, arrowheads, the dashed circle shows the lung), pancreatic neurons (**c**, the dashed circle shows the pancreas), and ENS cells (**c**, on the gut). d, Zoom-in images of various organ-intrinsic neurons as shown in (**a-c**). Scale bars: 1 mm (**a**), 100 μm (**b**, **c**), 50 μm (**d**).

**Extended Data Fig. 2:**
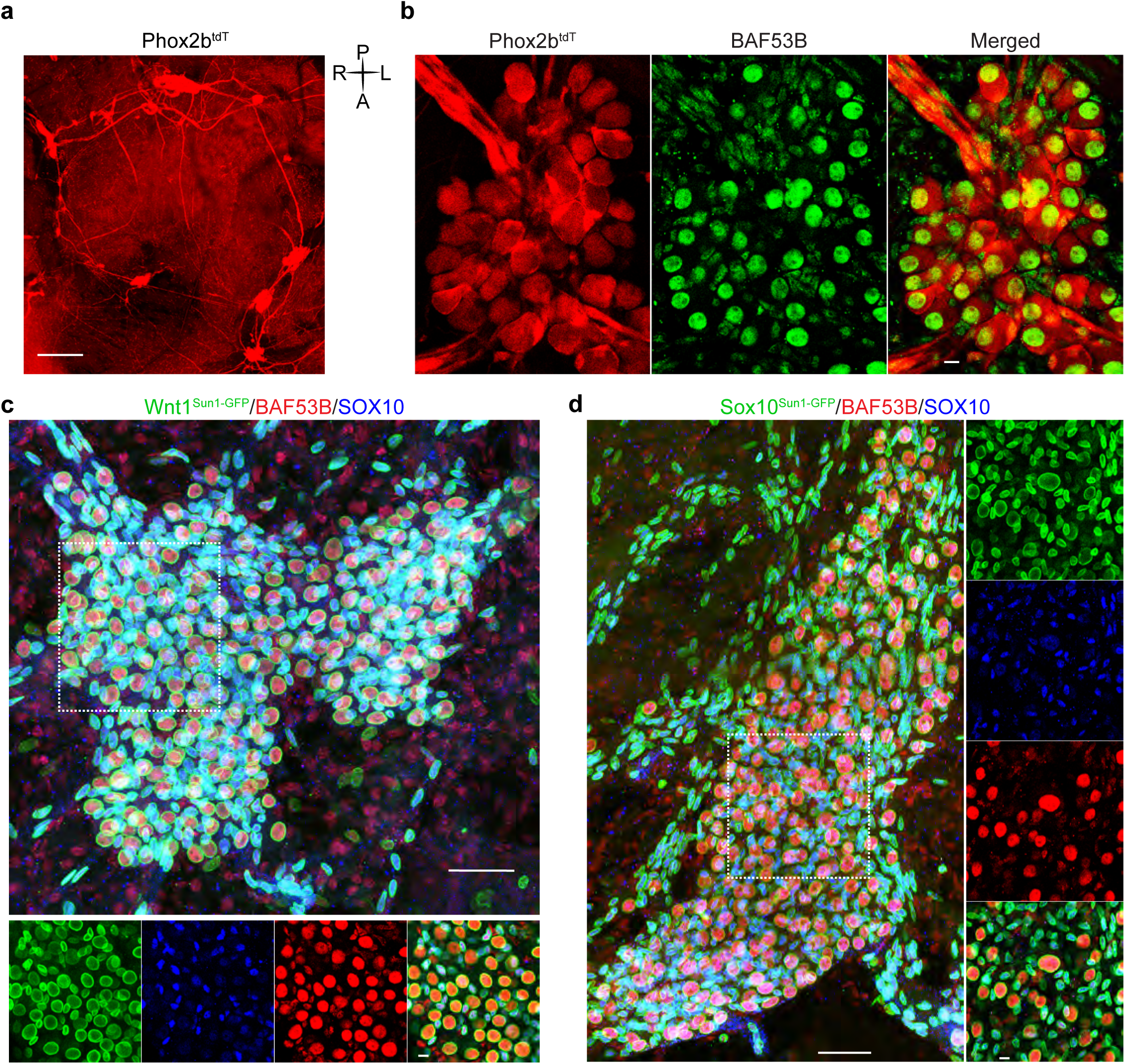
Lineage tracing of the ICNS. **a**, Superior view of a whole-mount heart from a one-month old Phox2b^tdT^ mouse showing the ICNS labelled with tdTomato (red). **b**, Representative ICNS ganglion from a Phox2b^tdT^ mouse heart co-labelled with tdTomato (red) and a pan-neuronal marker BAF53B (green), showing that all ICNS-neurons are derived from Phox2b lineage. c-d, Representative ICNS ganglia from Wnt1^Sun1-GFP^ (**c**) and Sox10^Sun1-GFP^ (**d**) hearts co-labeled with GFP (green, nuclear membrane), BAF53B (red), and SOX10 (blue), showing that all ICNS-neurons are derived from Wnt1 and Sox10 lineage. Zoom-in images from the dashed regions are shown at the bottom. Note that both Wnt1 and Sox10 lineages also mark glial cells and other heart cells. Scale bars: 500 μm (**a**), 10 μm (**b**, zoom-in for **c** and **d**), 50 μm (**c**, **d**)

**Extended Data Fig. 3:**
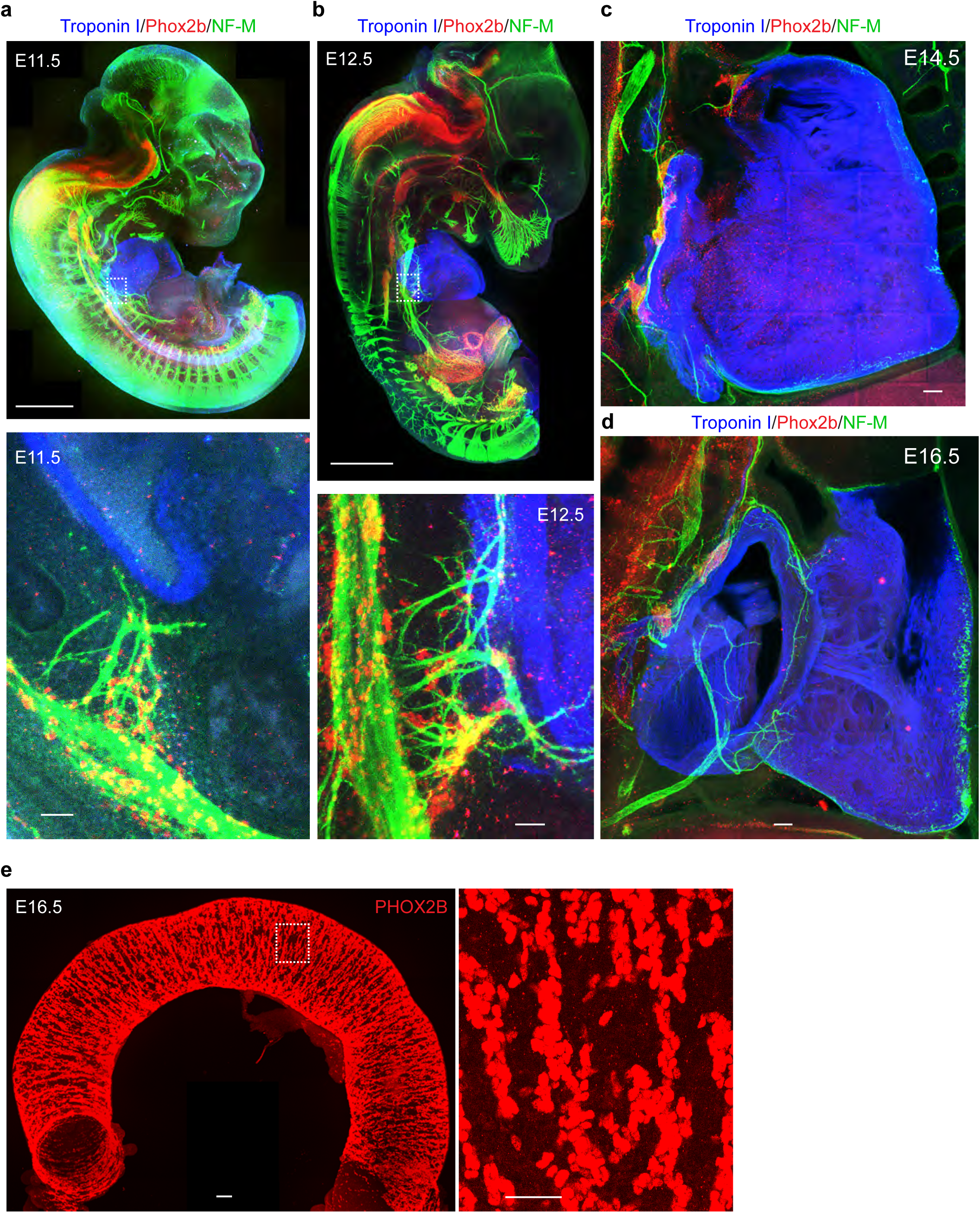
The ICNS lands on the heart at E12.5 via the vagus nerve. **a-d**, Whole-embryo 3D imaging of cleared mouse embryos at E11.5 (**a**), E12.5 (**b**), E14.5 (**c**), and E16.5 (**d**), stained with Phox2b (red), Neurofilament M (NF-M, green), and Troponin I (blue, heart), showing that ICNS cells (red) first land on the heart at E12.5 via vagal cardiac branches (green), with visible ganglion at E14.5, and discrete ganglia at E16.5. Zoom-in images from the dashed regions in (**a**) and (**b**) are shown at the bottom. **e**, Whole-organ 3D imaging of cleared mouse intestine at E16.5, stained with PHOX2B (red), showing extensive ENS cells colonizing the entire gut. Zoom-in images from the dashed regions (right) shows that clear ENS ganglia structures are absent at this age. Scale bars: 1 mm (**a**, **b**), 100 μm (**c-e**), 50 μm (zoom-in for **a**, **b**, and **e**).

**Extended Data Fig. 4:**
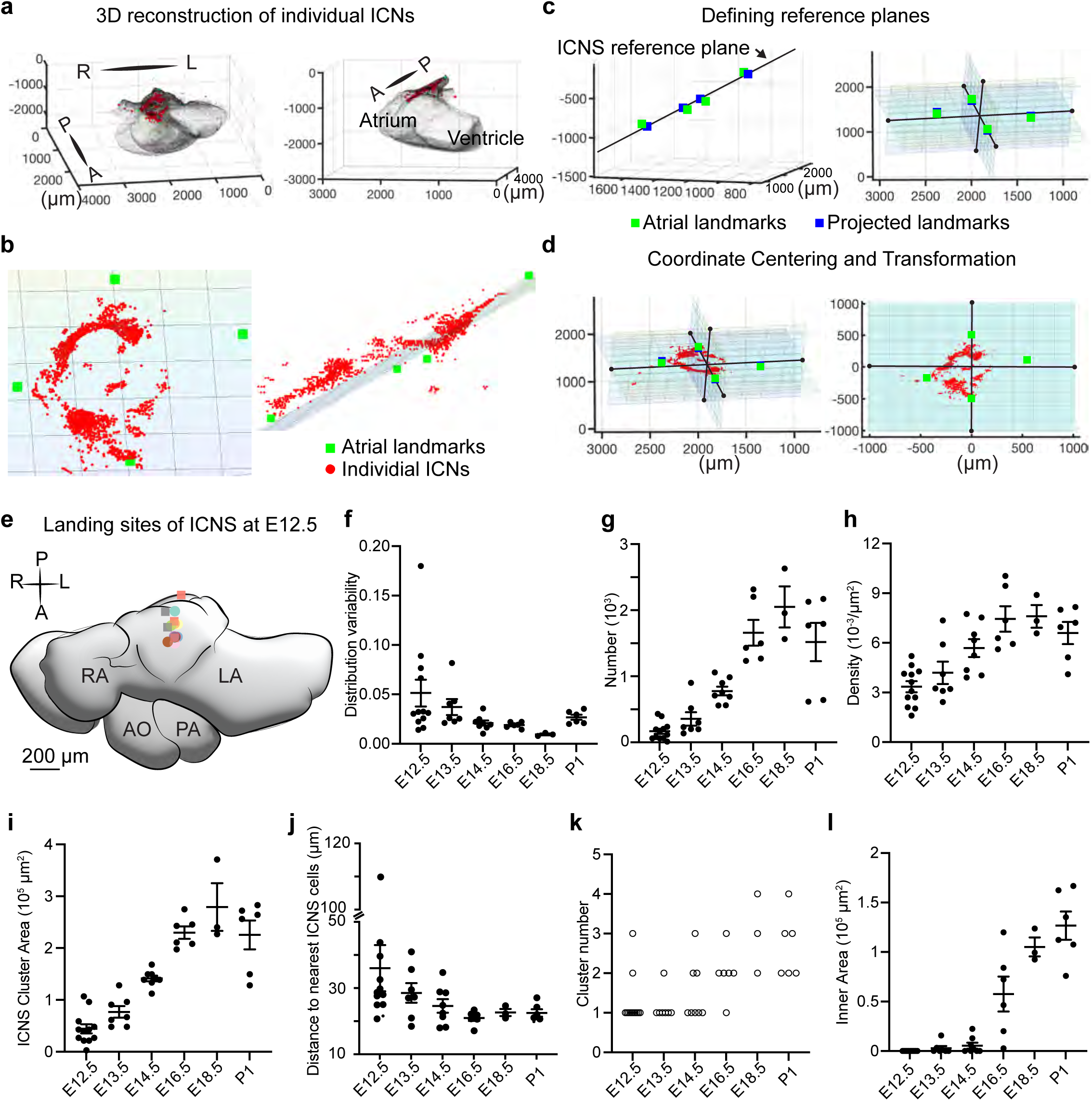
A computational approach to quantify the spatiotemporal distribution pattern of ICNS cells on the heart. **a**, 3D reconstruction of imaged ICNS cells (red, labelled with PHOX2B) on a wildtype mouse heart (grey, with background autofluorescence) using MATLAB. Superior (left) and left view (right) are shown. **b**, ICNS cells (red) as shown in (**a**) are largely located on a single plane (grey, with grids) defined by four consistent atrial landmarks (green square). **c**, Calculation of the ICNS reference plane defined by the four atrial landmarks (left) and generation of three corresponding axes (right) for re-orienting the heart across samples. Note that after this transformation, all heart samples are oriented in the same way on the new axes therefore spatial distribution can be compared and quantified across samples. **d**, Plots of ICNS cells (red) on the z = 0 plane, using axes before (left) and after transformation (right), showing the three reference planes. Note that after transformation, ICNS cells were predominantly located on the z = 0 plane and thus can be compared on the 2D plot. **e**, ICNS landing sites from 12 E12.5 mouse heart samples (left), superimposed on a schematic mouse E12.5 heart (superior view). Dots with different colors or shapes (circle, only one landing site; square, multiple landing sites) are from different samples. **f-l**, Quantifications of indicated parameters describing the spatiotemporal distribution patterns of the ICNS. mean ± sem.

**Extended Data Fig. 5:**
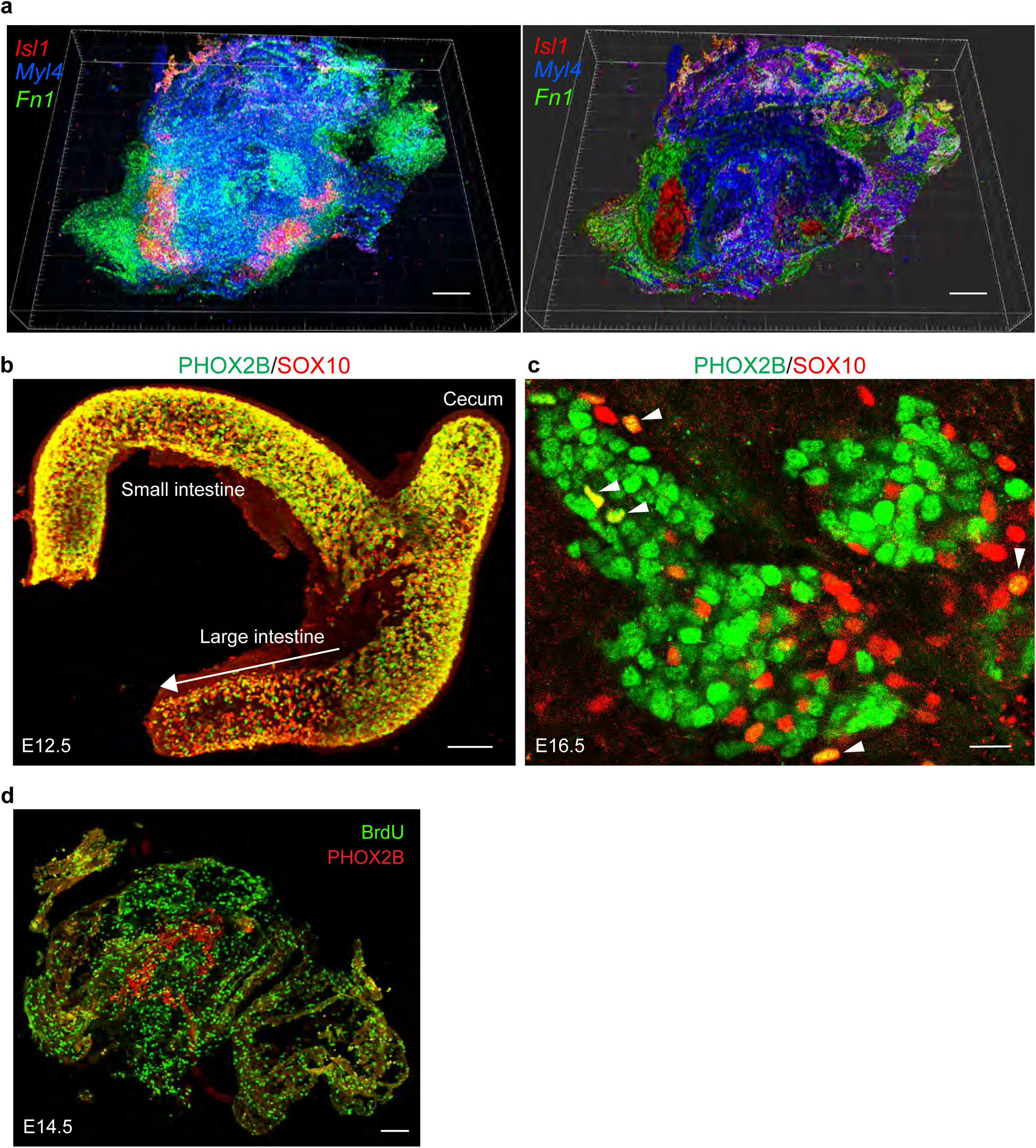
The cytoarchitecture around the ICNS. **a**, 3D reconstruction of 20 consecutive heart sections with RNAscope HiPlex analysis of indicated genes, showing in 3D views in Maximum Intensity Projection mode (left) and Normal Shading mode (right) by Imaris. Note that ICNS cells (*Isl1*^+^, red) are physically locked between cardiomyocytes (*Myl4*^+^, blue) and fibroblasts (*Fn1*^+^, green). **b**, Whole-organ 3D imaging of cleared mouse intestine at E12.5, stained with SOX10 (red) and PHOX2B (green), showing the migration wavefront (arrow, lack of PHOX2B signal). **c**, An E16.5 ICNS ganglion section, stained with SOX10 (red) and PHOX2B (green), showing the very sparse SOX10^+^/PHOX2B^+^ ICNS-precursors and no visible wavefront in the ICNS. **d**, Max projection of three ICNS-containing heart sections stained with PHOX2B (red) and BrdU (green), showing that both heart cells inside and outside of the ICNS ring are actively proliferating. Scale bars: 150 μm (**a**, **b**), 20 μm (**c**), 100 μm (**d**).

**Extended Data Fig. 6:**
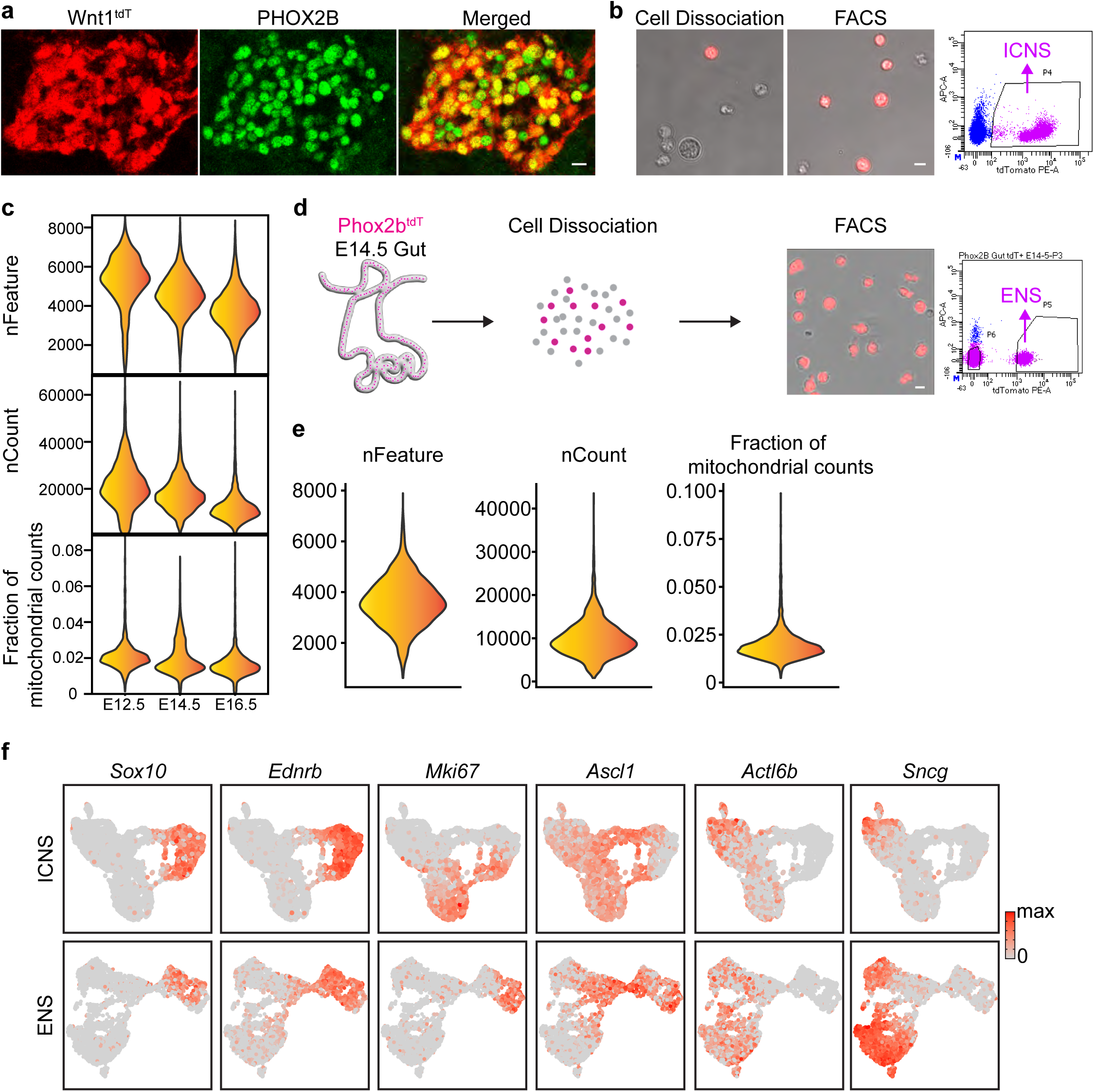
Single-cell transcriptomic profiling of ICNS cells at Phase I. **a**, ICNS ganglia from an E14.5 Wnt1^tdT^ mouse heart labelled with tdTomato (red) and PHOX2B (green) showing that ICNS cells are effectively labelled using Wnt1^tdT^ mice. **b**, Representative images of acutely isolated ICNS cells before (left) and after (middle) FACS purification. Cell selection parameters are shown on the right. **c**, Quality control of single-cell RNA-sequencing of E12.5 – E16.5 ICNS cells as shown in Fig. 2b. **d**, schematic illustration of single-cell RNA-sequencing of ENS cells acutely isolated and purified from the entire intestine of Phox2b^tdT^ mice. ENS cells after FACS purification and cell selection parameters are shown on the right. **e**, Quality control of single-cell RNA-sequencing of E14.5 ENS cells as shown in Fig. 2f. **f**, UMAP plots of ICNS and ENS cells as shown in Fig. 2e**-f**, coloured by expression of indicated cell state marker genes. Scale bars: 10 μm.

**Extended Data Fig. 7:**
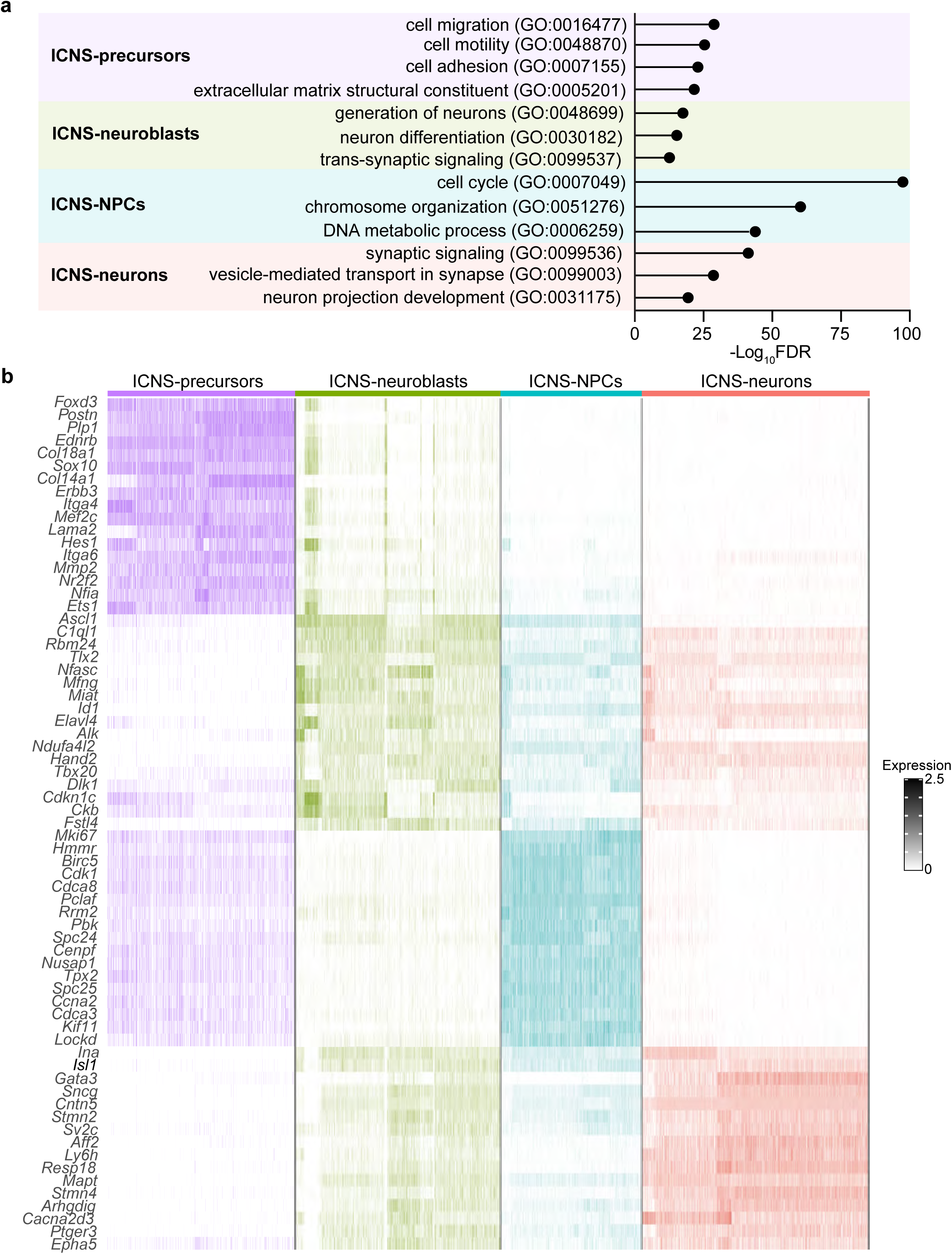
The four ICNS cell states. **a**, Top GO pathways of DEGs in indicated ICNS cell states. FDR, false discovery rate. **b**, Heat map of genes differentially expressed in indicated ICNS cell states.

**Extended Data Fig. 8:**
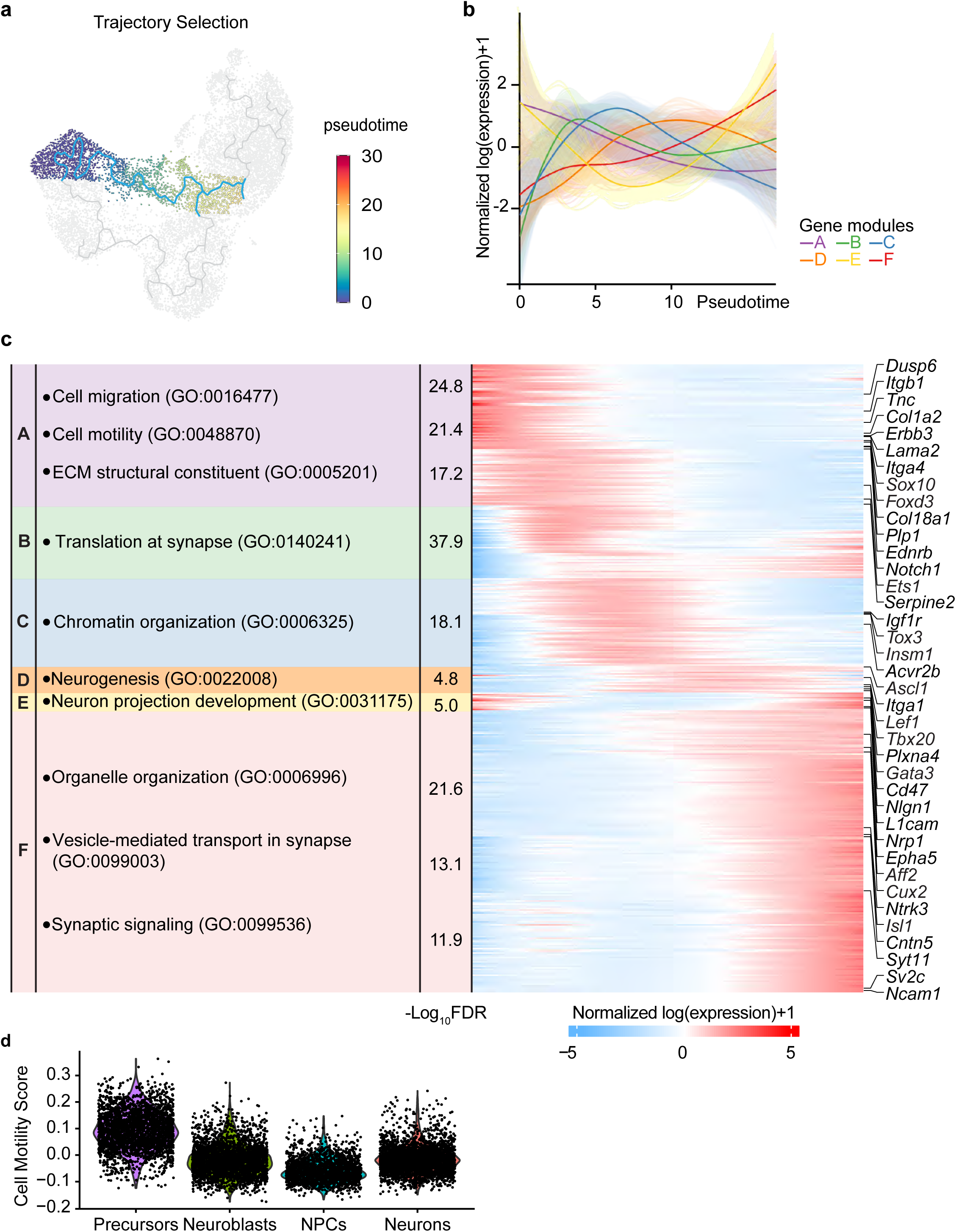
Trajectory analysis of the precursor-to-neuroblast path in the ICNS. **a**, Developmental trajectory from ICNS-precursors to ICNS-neuroblasts predicted using Monocle 3. **b**, Expression level of six gene modules (color coded) identified using hierarchical clustering along the predicted pseudotime. **c**, Selected Top GO pathways of genes in indicated gene module (left) with heatmap showing gene expression patterns along the predicted pseudotime (right), showing genes involved in cell migration and motility are immediately turned off during differentiation. FDR, false discovery rate. **d**, Cell motility score of various ICNS cell states, showing a significant reduction of migratory motility after ICNS differentiation.

**Extended Data Fig. 9:**
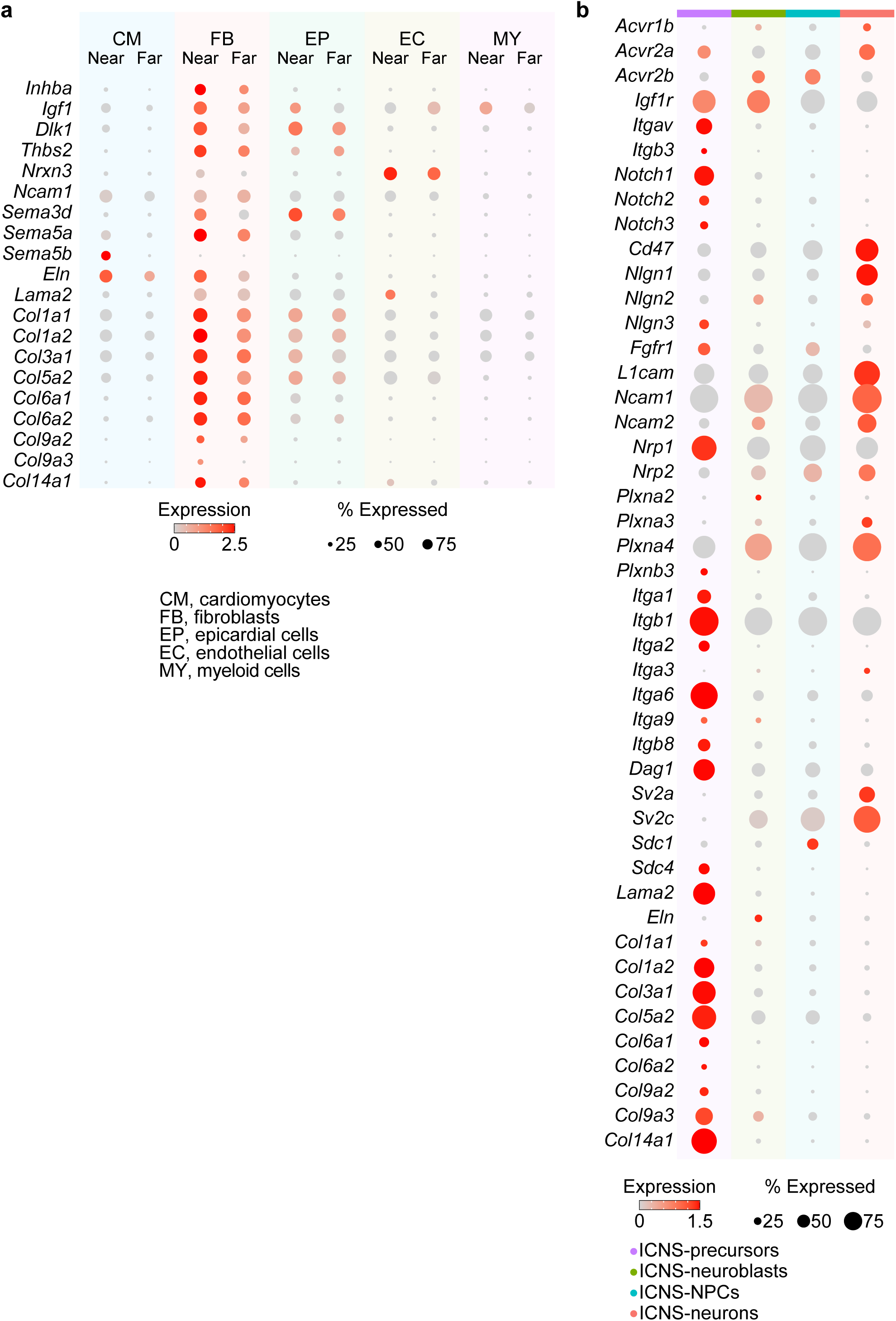
Genes differentially expressed in heart^near^, heart^far^, and ICNS cells. **a-b**, Dot plots of indicated ligands between heart^near^ and heart^far^ cells separated by cell types (**a**) and corresponding receptors/ECM genes between various ICNS cell states (**b**).

**Extended Data Fig. 10:**
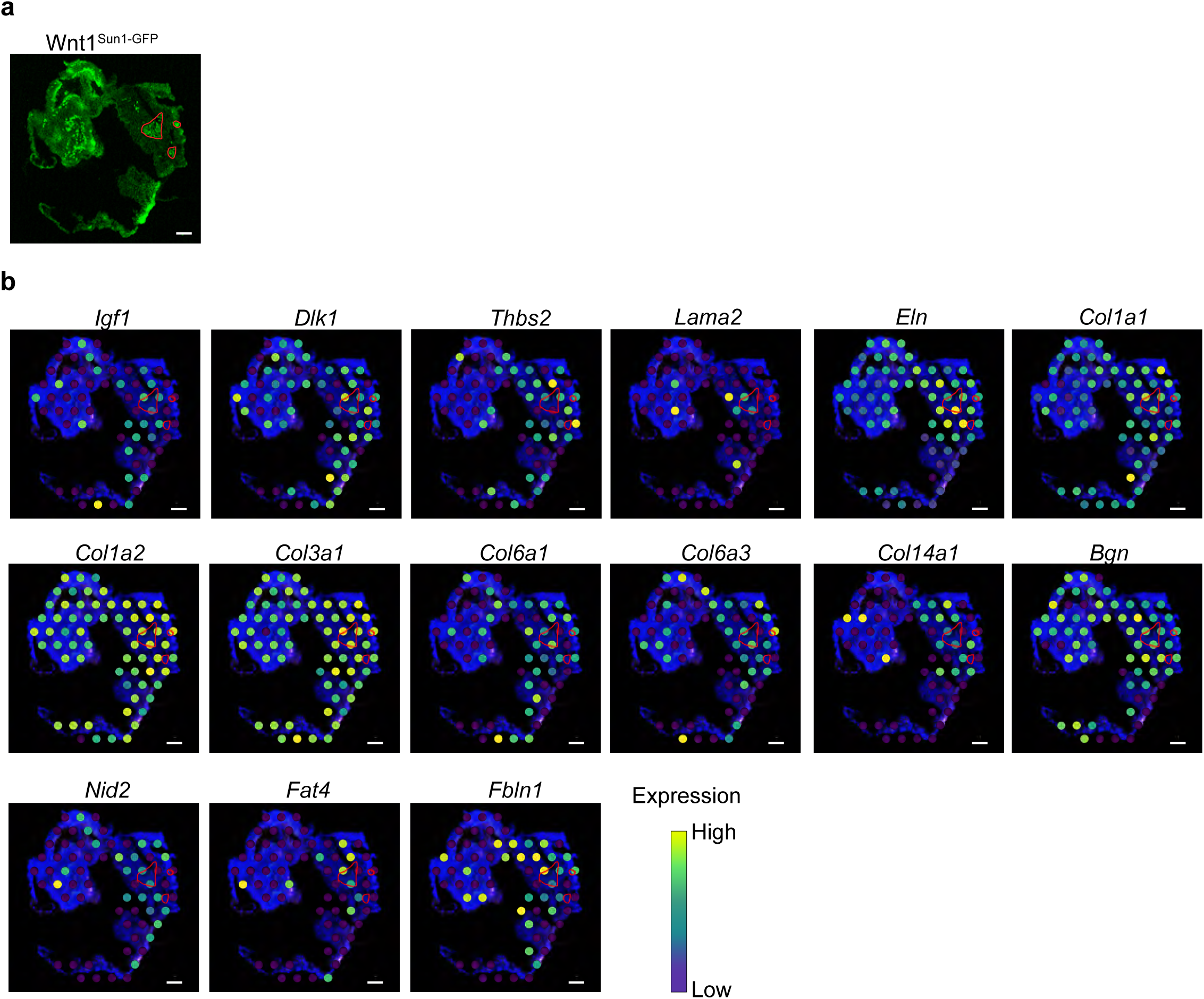
Spatial transcriptomics of the heart containing the ICNS. **a**, An E14.5 Wnt1^Sun1-GFP^ heart section with native GFP signal (green) showing the location of the ICNS (red circles). **b**, 10x Visium based spatial transcriptomics barcoded spots of the section as in (a), coloured by expression of indicated genes that are enriched in the ICNS area. Scale bar: 100 μm.

**Extended Data Fig. 11:**
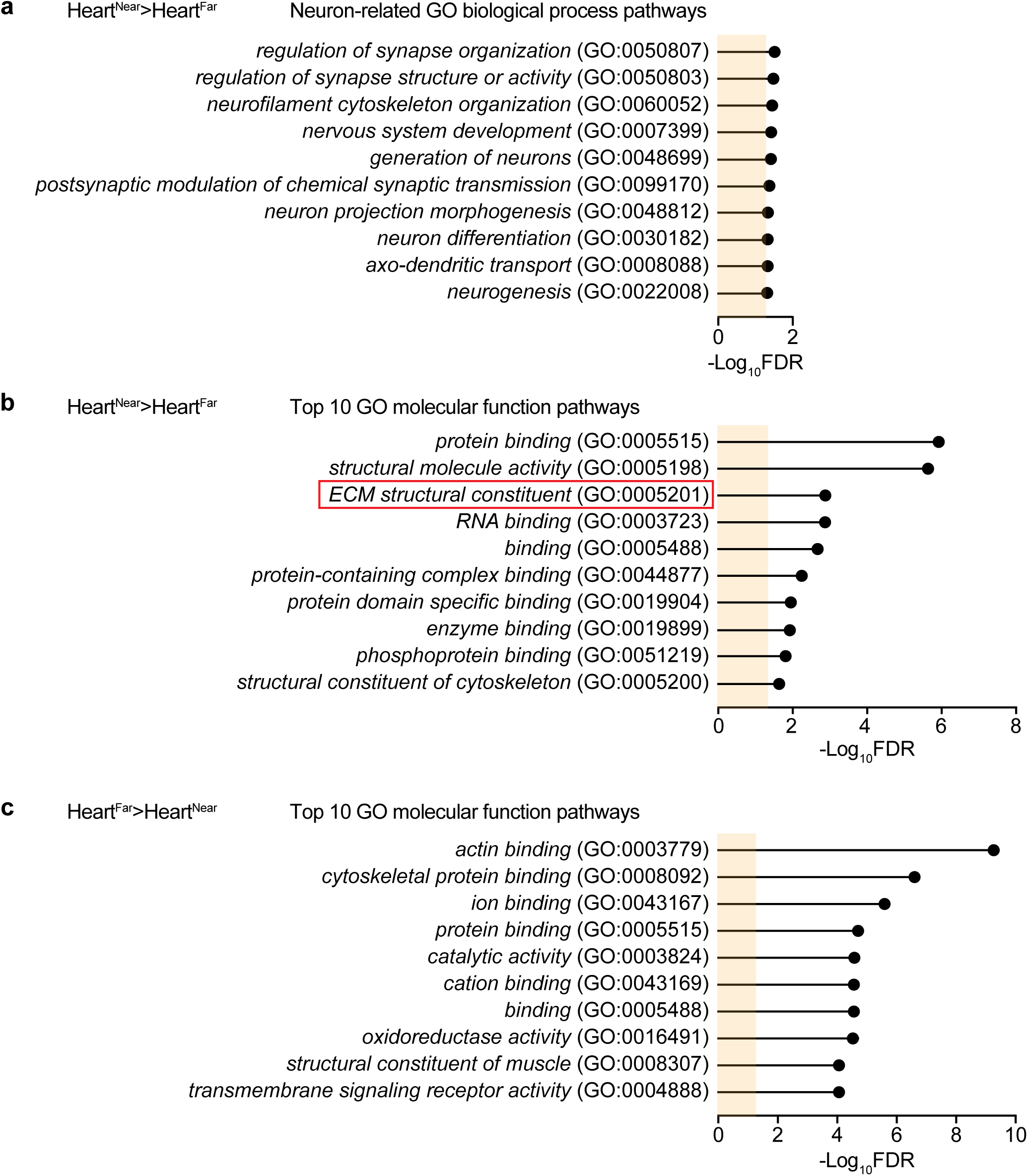
GO pathway analysis of proteins enriched around the ICNS area on the heart. **a**, Selected top GO biological process pathways of proteins enriched in heart^near^ over heart^far^. b-c, Top 10 GO molecular function pathways of proteins enriched in heart^near^ over heart^far^ (**b**) or in heart^far^ over heart^near^ (**c**). FDR, false discovery rate. Yellow shades indicate FDR = 0.05.

**Extended Data Fig. 12:**
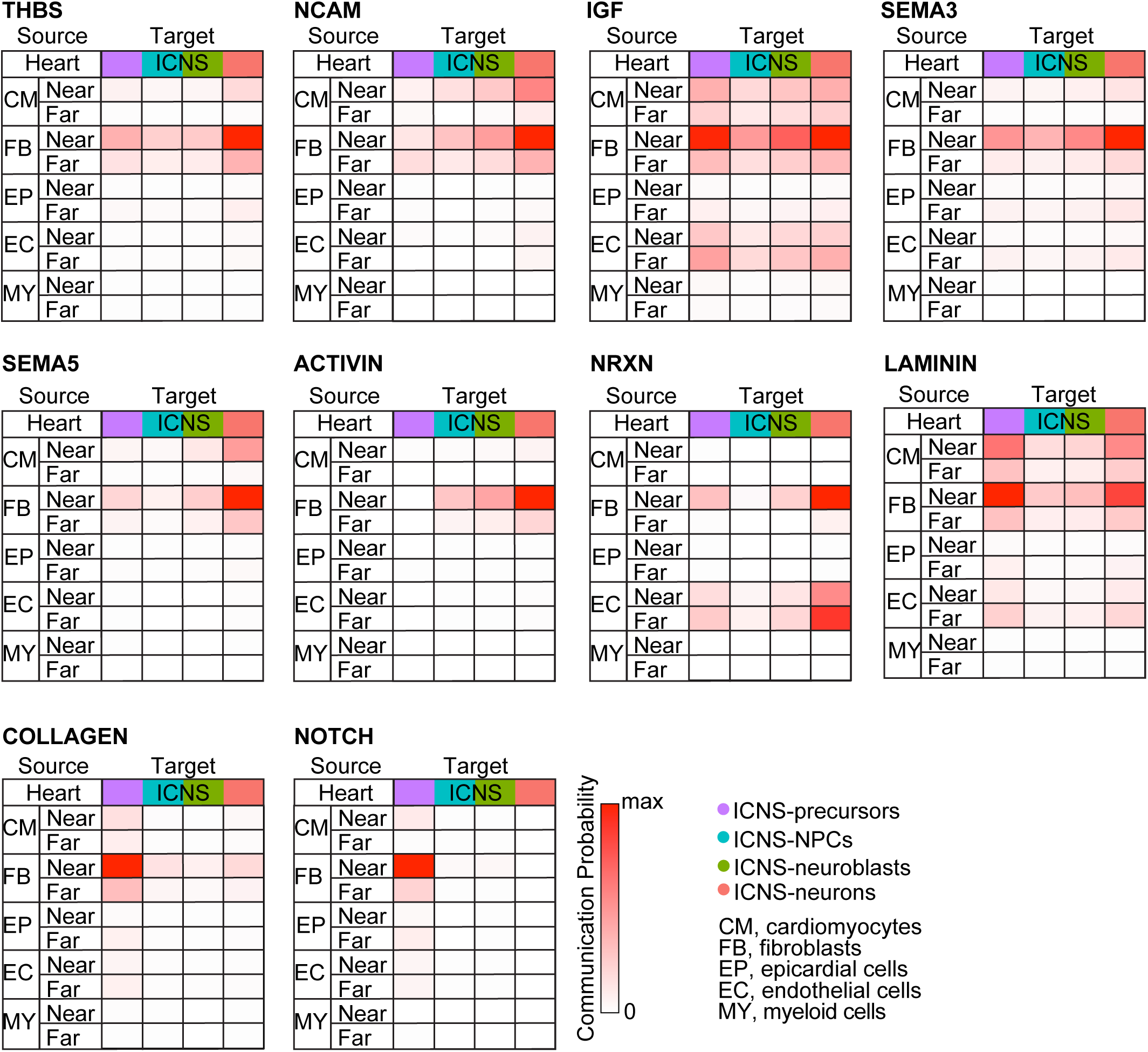
Cell-cell communication analysis between heart and ICNS cells. Heatmaps of communication probabilities from CellChat analysis using indicated cell types from E14.5 heart^near^ (near) and heart^far^ (far) single-cell RNA-sequencing datasets as shown in Fig. 3b as donors (source) and indicated ICNS cell states at E14.5 (color coded) as recipients (target). CM, cardiomyocytes; FB, fibroblasts; EP, epicardial cells; EC, endothelial cells; MY, myeloid cells.

**Extended Data Fig. 13:**
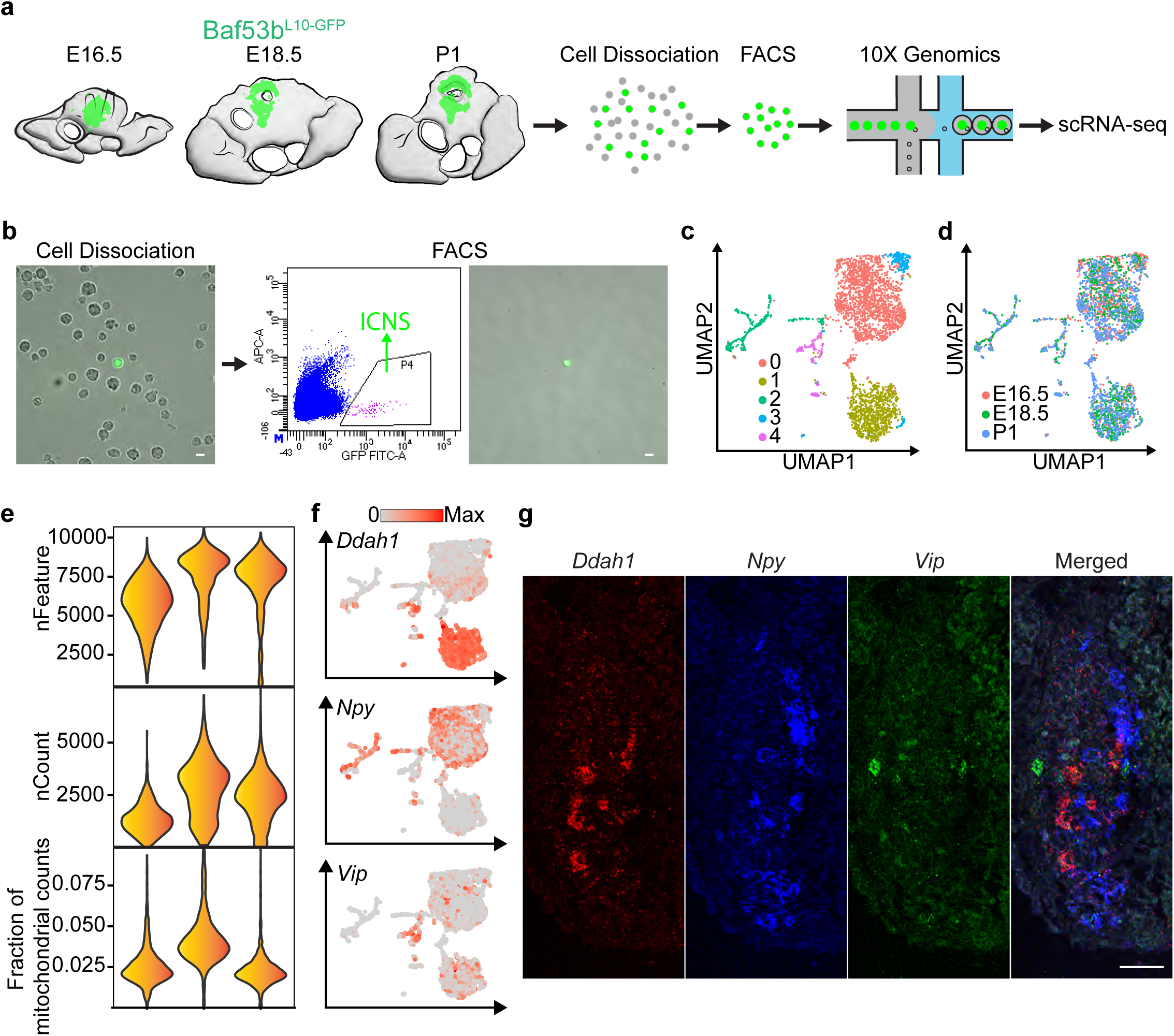
Single-cell transcriptomic profiling of maturing ICNS cells at Phase II. **a**, Schematic illustration of single-cell RNA-sequencing of fluorescently labelled ICNS cells acutely isolated and purified from E16.5 – P1 Baf53b^L10-GFP^ mouse hearts. **b**, Representative images of acutely isolated ICNS cells before (left) and after (middle) FACS purification. Cell selection parameters are shown on the right. **c-d**, UMAP plots of integrated E16.5 – P1 ICNS-neuron datasets, colored by genetically defined Seurat clusters (**c**) and age (**d**). **e**, Quality control of single-cell RNA-sequencing of E16.5 – P1 ICNS-neurons as in (**c**). **f**, UMAP plots of ICNS-neurons as in **(c)**, coloured by expression of indicated cell type marker genes. **g**, RNAscope HiPlex analysis of indicated genes in an ICNS ganglion, showing different ICNS-neuron subtypes. Scale bars: 10 μm (**b**), 50 μm (**g**).

**Extended Data Fig. 14:**
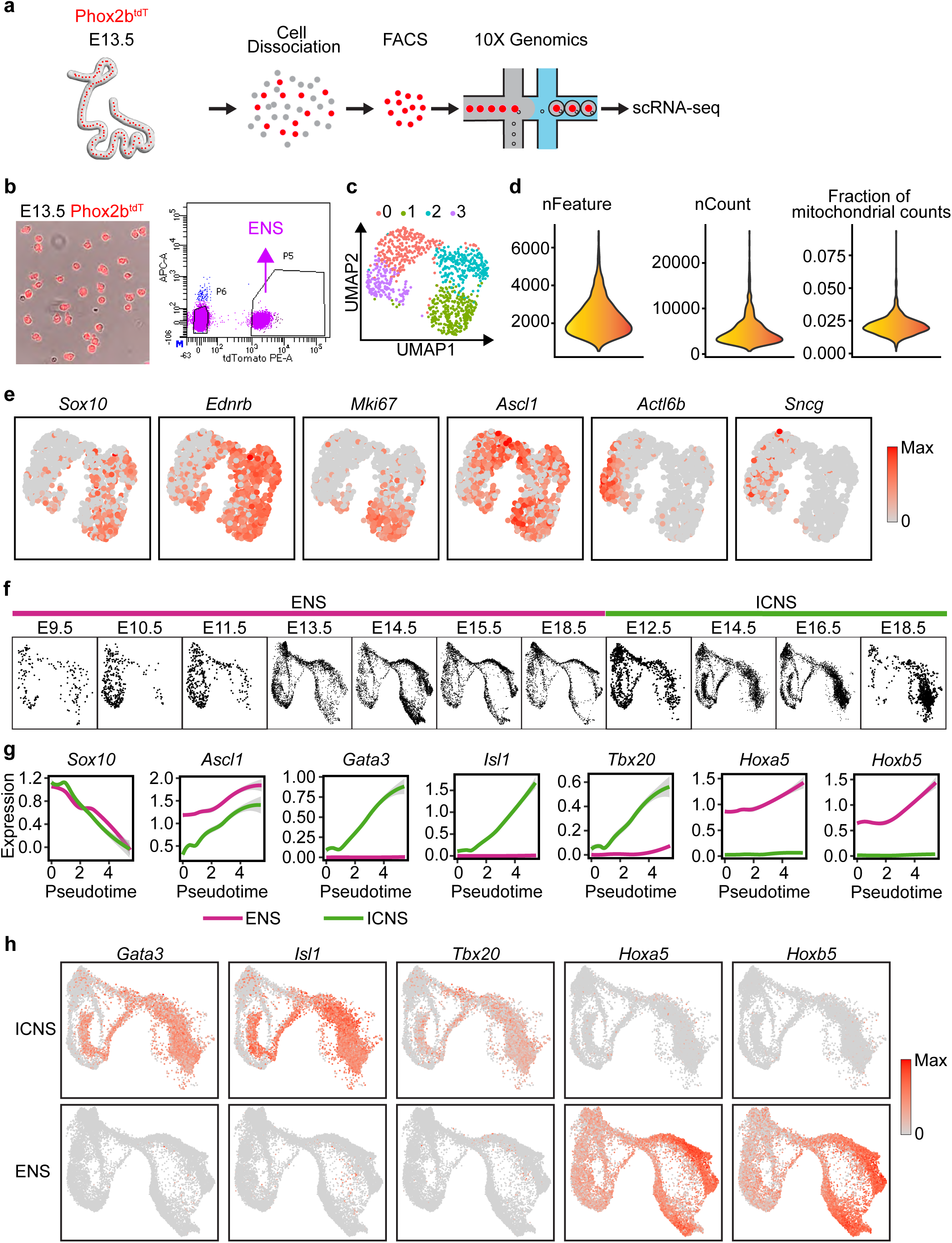
Comparisons between single-cell transcriptomics of developing ICNS and ENS cells. **a**, Schematic illustration of single-cell RNA-sequencing of fluorescently labelled ENS cells acutely isolated and purified from E13.5 and E14.5 Phox2b^tdT^ mouse intestines. **b**, Representative images of acutely isolated and FACS-purified E13.5 ENS cells (left) and cell selection parameters (right). Scale bar: 10 μm. **c**, UMAP plots of E13.5 ENS cells, colored by genetically defined Seurat clusters. **d**, Quality control of single-cell RNA-sequencing of E13.5 ENS cells as in (**c**). **e**, UMAP plots of ENS cells as in **(c)**, coloured by expression of indicated cell state marker genes. **f**, UMAP plots of integrated ENS and ICNS cells across development as shown in Fig. 4a, separated by age as indicated. **g**, Expression of indicated genes, calculated as log (counts/size factor + 1), in ICNS (green) and ENS (magenta) precursor/neuroblast cells along the predicted pseudotime as shown in Fig. 4g. Fitted by loess (locally estimated scatterplot smoothing), mean ± 95 % ci. **h**, UMAP plots of integrated ENS and ICNS cells as shown in Fig. 4a, coloured by expression of indicated ICNS- and ENS-specific transcription factor genes. FDR, false discovery rate.

**Extended Data Fig. 15:**
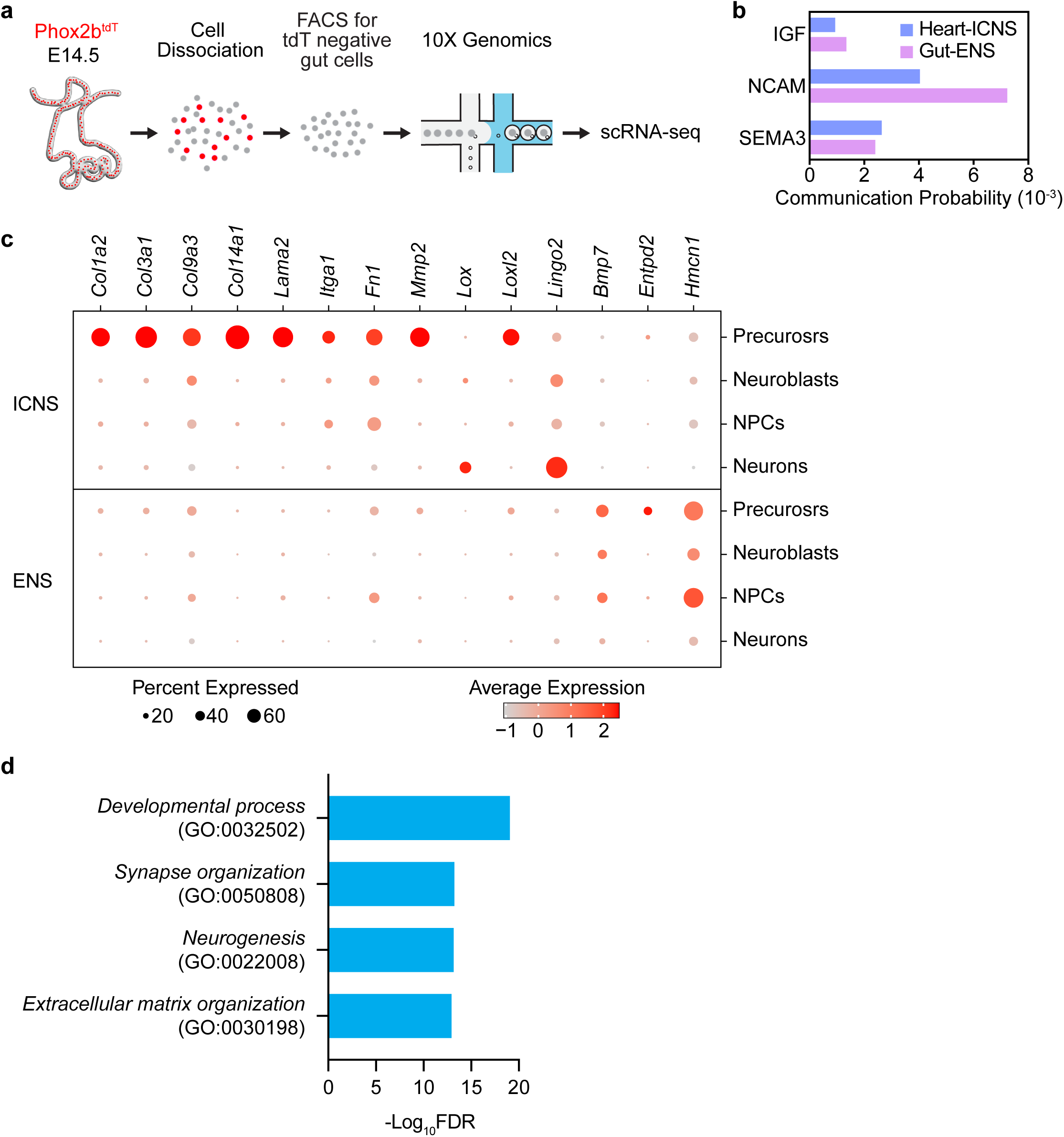
Cell-cell interaction analysis between visceral organs and their intrinsic neurons. **a**, Schematic illustration of single-cell RNA-sequencing of fluorescence negative ENS-free gut cells acutely isolated and purified from E14.5 Phox2b^tdT^ mouse intestines. **b**, Communication probability of indicated neuronal maturation pathways in heart-ICNS and gut-ENS donor-recipient pairs. **c**, Dot plots of indicated ECM genes differentially expressed ICNS (top) and ENS (bottom) in indicated cell states. **d**, Selected top GO biological process pathways of DEGs enriched in transcriptionally shifted ENS^heart^ precursors as shown within the dashed circle in Fig. 5f.

**Supplementary Table 1: Detailed information for anatomical imaging and single-cell RNA-sequencing.** Extended Data Table. 1 describes sample processing methods and age. Extended Data Table. 2 describes antibodies and dilutions for all imaging figures. Extended Data Table. 3 describes a summary of antibodies used in this manuscript and their sources/catalogue numbers. Extended Data Table. 4 describes details for all single-cell RNA-sequencing experiments, including mouse genotypes, ages, number of animals used, tissues collected, quality control standards.

## Supplementary Video Legends

**Supplementary Video 1: Simulation of the ICNS organization during surrounding heart cell proliferation.** Red, ICNS cells; green, heart cells in the ICNS inner area; blue, heart cells outside the ICNS ring. Each simulation step is a frame in the video. Video related to Fig. 1h.

## Methods

### Animals

All animal husbandry and procedures were performed in compliance with Yale University’s Institutional Animal Care and Use Committee and National Institute of Health (NIH) guidelines. Both male and female animals were used for the experiments and no differences between sexes were observed.

#### Mouse lines

Wild-type C57BL/6J (000664), Wnt1-cre (022501), Sox10-cre (025807), Baf53b-cre (027826), Phox2b-Flpo (022407), lox-tdTomato (007914), frt-tdTomato (032864), lox-Sun1-sfGFP^25^ (021039) were from the Jackson Laboratory. lox-L10-eGFP mice were described before^52^.

### Single-cell RNA-sequencing and data analysis

#### Cell dissociation and sequencing

Heart and gut (entire intestine) tissues were dissected out from indicated mice into ice-cold Leibovitz’s L-15 Medium (Gibco^TM^). The heart region that contains the ICNS was further identified and trimmed using a Leica M205FCA Fluorescence Stereo Microscope. For heart^far^ cells (Fig. 3a), a piece of heart region that does not contain the ICNS was dissected for sequencing. Dissected tissues were cut into 1-3 mm^2^ pieces and incubated with 0.25% trypsin-EDTA (Gibco^TM^) at 37 °C for 10 minutes (E12.5 – E14.5) or 20 minutes (E16.5 – P1). The tissues were then washed with L-15 containing 10% fatal bovine serum (FBS) and incubated with 2 mg/mL Collagenase A and 2 mg/mL of Dispase II (in 1 x HBSS) at 37°C for 30 minutes (E12.5 – E14.5) or 40 minutes (E16.5 – P1). After washing (L-15 with 10% FBS), cells were manually triturated using three fire-polished Pasteur pipettes with decreasing opening sizes. Maturing ICNS-neurons isolated from Baf53b^L10-GFP^ mice at E16.5, E18.5, and P1 were further purified using 30% and 60% Percoll density gradient. The cell suspension was filtered through a 40 μm cell strainer (Corning). Cells were centrifuged at 200 x g for 10 minutes at 4°C and re-suspended in 1x HBSS containing 0.04% BSA. Fluorescently labelled cells were sorted on a BD FACS Aria cell sorter and collected in ice-cold 1x HBSS containing 0.04% BSA at Yale Flow Cytometry Facility. Approximately 5,000-10,000 cells were loaded in each channel of the 10x microfluidic device to target for 3,000-6,000 cells as an output of one sample. Single-cell cDNA libraries were prepared with the Chromium Single Cell 3’ V3 reagent kit at the Yale Center for Genomic Analysis (YCGA) and sequenced using an Illumina NovaSeq S4 sequencer at 150-300 million reads to achieve a fine sequencing depth of 30,000-50,000 reads per cell.

#### Basic bioinformatic processing

Transcriptomic data were aligned to the mm10-2020-A mouse genome reference using the Cell Ranger software v.7.1.0 (10X Genomics). The following quality control metrics were applied to filter low-quality cells: number of genes per cell > 200; number of genes per cell < 20000; percentage of mitochondria genes < 10%. Single-cell RNA-sequencing datasets were integrated (if indicated) and processed using the R package Seurat v.4.3.0^31^. Cell clusters were identified using top 50 principal components visualized using UMAP^32^. ICNS and ENS cells states (progenitors, neuroblasts, NPCs, and maturing neurons, such as in Fig. 2e-f) as well as different heart and gut cell types (such as in Fig. 3c) were selected based on Seurat clusters expressing indicated corresponding marker genes. Cell cycle analysis (Fig. 2i) was performed using the CellCycleScoring function implemented in Seurat. DEGs between indicated cell groups (such as in Extended Data Fig. 9) were identified using the bimod likelihood-ratio test implemented in Seurat. GO pathway analysis of DEGs (such as in Fig. 2l) were performed using the Gene Ontology Resource GO Enrichment Analysis tool^36,37^ (http://geneontology.org).

Genes within indicated GO terms were identified using the Mouse Genome Informatics (MGI) database (https://www.informatics.jax.org/vocab/gene_ontology). Mouse transcription factors were identified using the AnimalTFDB 3.0 mouse database^53^ (http://bioinfo.life.hust.edu.cn/AnimalTFDB/#!/). The AddModuleScore^54^ function implemented in Seurat was used to calculate cell motility score (Extended Data Fig. 8d, using genes within GO:0048870). Similarity of co-cultured ENS cells to ICNS or ENS cells was calculated separated, using DEGs enriched in ICNS or ENS cells. For precursor similarity score shown in Fig. 5i, top 200 DEGs enriched in ICNS precursors and top 200 DEGs enriched in ENS precursors were used respectively. For ECM similarity score shown in Fig. 5k, all DEGs between ICNS and ENS precursors within ECM GO: 0031012 were used (80 for ICNS, 15 for ENS). ICNS-ENS precursor similarity score (Fig. 5i) and ICNS-ENS ECM similarity score (Fig. 5k) were calculated by subtracting the ENS similarity score from the ICNS similarity score.

#### RNA velocity inference

We estimated the cell transition of ICNS cells using RNA velocity, focusing on the E14.5 dataset (Fig. 2h). We first used *velocyto run10X* function^55^ to obtain the spliced and unspliced read counts matrix. Then we inferred the RNA velocity by following the scVelo (v0.2.5) pipeline. We first kept the genes with at least 20 of cells expressed in both unspliced and spliced matrices and then selected the top 2000 highly variable genes based on dispersion to construct the PCA space. Next, we computed moments for velocity estimation for each cell across its 30 nearest neighbours, where the neighbour graph was calculated based on the top 30 PCs. We then calculated the velocity using function *tl.velocity()* with mode set as “stochastic” and *tl.velocity_graph()*, with all parameters set as default and estimated the velocity pseudotime using function *tl.velocity_pseudotime()*. We visualized our results using *pl.velocity_embedding_stream()*.

#### Trajectory inference

Monocle3^38^ v1.3.1 was used to infer the trajectory of the integrated ICNS data (Extended Data Fig. 8) and the ICNS-ENS integrated data (Fig. 4). The beginning of the trajectory was determined by 20-50 cells with the highest *Sox10* expression in the precursor state. The branch in Extended Data Fig. 8 was selected by the “choose_cells” function. We first utilized the “graph_test” function with q_value < 0.01 to isolate the variable genes along the target branch. Then we computed the gene expressions over pseudotime to create a time series of gene expressions. Then we further applied an expression range cut-off across the pseudotime to retain highly variable genes. We next performed hierarchical clustering^39^ and classified the genes into groups based on their expressions over the pseudotime using the “cutree“ function with a cluster number set at 13 (k=13). Within the 13 there are groups sharing the same expression patterns, therefore we further combined the groups with shared expression patterns and generated the final 6 distinct modules shown in Extended Data Fig. 8. The GO analysis of the module genes was performed using the Gene Ontology Resource GO Enrichment Analysis tool^36,37^ (http://geneontology.org).

#### scTIE framework

The integrated ICNS and ENS dataset (ICNS, E12.5 – E18.5; ENS, E13.5 – E18.5) was used for scTIE analysis. As the scTIE was originally designed for paired single-cell RNA-sequencing and ATAC-seq data^48^, we first modified scTIE for single-cell RNA-sequencing data by removing the encoder and decoder for ATAC-seq and the modality alignment loss. We used the integrated matrix of 2000 highly variable genes returned by Seurat as input. scTIE is designed to model the cell transition between two developmental stages using iterative optimal transport (OT). To model the cell state transitions along the differentiation trajectory, we divided both the ICNS- and ENS-cell trajectories into two stages, early (mainly include ICNS- and ENS-precursor and neuroblast cells) and late (mainly include ICNS- and ENS-neuron cells). We used the estimated pseudotime inferred by monocle to decide the cut-off, which is set as 5.5 for ENS-cell trajectory and 6.5 for ICNS-cell trajectory. We then ran the modified version of scTIE by setting the weights of the reconstruction loss as 10, the weights of OT as 0.1, the number of epochs as 500 with OT updated every 10 epochs. Next, we used the estimated transition probability matrix from scTIE 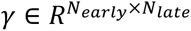 to calculate the cell type transition probability, where *N_early_* and *N_late_* indicates the number of cells at the early and late stage respectively. *γ*(*i*, *j*) can be considered as the probability of cell *i* at the early stage transits to cell *j* at the late stage. We then summarized this to calculate the cell fate transition to the ICNS and ENS. For each cell *i* at the early stage and *G_ICNS_* and *G_ENS_* at the late stage, the transition probabilities of ICNS 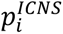 and ENS 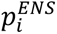 were then calculated as the following.

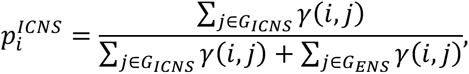

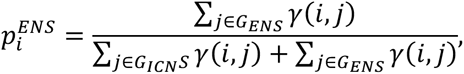

where *G_early_* is the set of all the cells at the early stage, *G_ICNS_* contains the ICNS-cells at the late stage and *G_ENS_* contains the ENS-cells at the late stage. We then predicted the cell fate transition probabilities of the cells at the early stage by finetuning a one-layer classifier on the pretrained features in the embedding space, with the L1 regularization weights set as 0.1, and backpropagate the gradients from the prediction layer to select the top genes that are predictive to the cell transition.

#### Cell-cell communication inference

To calculate cell-cell interactions between visceral organ cells and their organ-intrinsic neurons (Figs. 3e, 3i, 5a-d, Extended Data Figs. 12 and 15b), we merged the single-cell RNA-sequencing Seurat objects of the organ cells (heart^near^, heart^far,^ and Gut at E14.5) with organ-intrinsic neuron datasets (E12.5 – E18.5 for ICNS and E9.5 – E18.5 for ENS). For the CellChat analysis of the cocultured experiments, we merged the two single-cell RNA-sequencing datasets of the heart-ENS cocultures and the four single-cell RNA-sequencing datasets of the gut-ENS cocultures (Fig. 5k). We transformed the merged Seurat objects into the CellChat objects using the CellChat^56^ package v1.6.1. We incorporated the CellChat object with the “CellChatDB.mouse” database, then utilized the CellChat functions to identify overexpressed genes and interactions and computed their communication probabilities. We utilized the “subsetCommunication” function to obtain the ligand-receptor pairs with p-value < 0.05 and specified the organ cells as “source” and organ-intrinsic neuron cells as “target”. “Secreted Signaling”, “Cell-Cell Contact” and “ECM-Receptor” are the three signalling annotations in the CellChatDB.mouse database. In each signalling annotation, there are different pathways. We extracted and summed the communication probabilities of the ligand-receptor pairs to acquire the total communication probabilities of each signalling annotation or different pathways. Heat maps showing the interaction strength of pathways (Extended Data Fig. 12) were plotted using the GraphPad Prism. The “netVisual_circle” function built in CellChat was used to plot Fig. 3e.

### Spatial transcriptomics with 10x Visium

Hearts from Wnt1^sun1-GFP^ E14.5 mice were acutely dissected and freshly frozen in cryo-embedding media (OCT). Cryosections (10 μm) were cut using a cryostat (Thermo Fisher Scientific) and mounted onto the 10x Visium Spatial Gene Expression Slide. Tissue sections were imaged with a Leica SP8 confocal microscope to obtain tissue morphology and locate ICNS ganglia (Extended Data Fig. 10a). Sequencing libraries were prepared following the manufacturer’s protocol (10x Visium) and sequenced using an Illumina NovaSeq S4 sequencer at YCGA. Raw FASTQ files and corresponding confocal images were processed by the Space Ranger software 2.0.0 (10x Genomics). We pre-processed the 10x Visium dataset using log-normalization^57,58^. Gene expression values for each spot were normalized to the total number of transcripts and multiplied by a scaling factor of 10,000. The normalized dataset was then transformed to log scale using log1p. The log-normalization process was performed using the Python package Scanpy 1.8.2^58^. Following this standard data pre-processing, we visualized gene expression levels within spots on a high-resolution histology image to examine spatial distributions of genes. The high-resolution image is a down-sampled version of the original full-resolution histology image, typically generated by the Space Ranger pipeline. It has 2,000 pixels in its largest dimension. To map spatial spots onto the high-resolution image, we utilized a text file that describes spot coordinates and a JSON file that describes image properties. The text file and the JSON files are outputs from the Space Ranger pipeline. The text file contains the pixel coordinates of spot centers on the full-resolution image, while the JSON file includes the spot diameter in pixels on the full-resolution image, as well as a scaling factor that transforms pixel positions from the original full-resolution image to the high-resolution image. Additionally, we visualized region of interest (ROIs) for the ICNS on the high-resolution histology image. We circled and saved ROIs using the Fiji software and loaded ROIs into Python using Python module read-roi 1.6.0 for the joint visualization of ROIs, gene expression levels within spots, and the high-resolution histology image.

### RNAscope HiPlex assay

RNAscope HiPlex assays were performed following the manufacturer’s protocol (Advanced Cell Diagnostics) as previously described^59^. The E16.5 mouse heart and gut were acutely dissected and freshly frozen in cryo-embedding medium (OCT). Cryosections (10 μm) were cut using a cryostat (Thermo Fisher Scientific), mounted onto Superfrost Plus slides (Thermo Fisher Scientific), and stored at −80 °C until use. Slides were immediately immersed into fresh 4% paraformaldehyde (PFA) in RNase-free PBS for 60 minutes at room temperature, followed by dehydration with 50%, 70% and 100% ethanol. Samples were then digested with Protease IV for 30 minutes at room temperature. After hybridization with designed probes (see Supplementary Table 1) for 2 hours at 40 °C, the sections were treated with HiPlex Amp and then HiPlex Fluoro solutions for the target group. Slides were not mounted after HiPlex Fluoro and instead imaged in 4xSSC using a Leica SP8 confocal microscope equipped with a motorized stage, a PMT detector, two HyD SP detectors, and four laser lines (405 nm, 488 nm, 552 nm and 638 nm) and a 16× immersion objective (HC FLUOTAR L 16×/0.8 IMM motCORR VISIR). After each group, the fluorophores were cleaved by 10% cleaving solution (ACD, Cat# 324130) and then hybridized with solutions for the next group. After each round imaging (4 groups/12 probes), probes were removed using the HiPlexUp reagent, and sections were hybridized with another 12 probes (2 hours, 40°C) for the next round analysis. RNAscope HiPlex Assays for the following genes were performed (Figs. 1g, 3f, and 4c, Extended Data Fig. 13g): *Ddah1* (R1T4), *Hoxa5* (T2T4), *Vip* (R1T6), *Phox2b* (R3T6), *Npy* (R1T7), *Isl1* (R2T7), *Fn1* (R2T8), *Wt1* (R3T8), *Lox* (R1T9), *Myl4* (R3T10), *Dcn* (R3T11), *Pecam* (T3T12).

### Whole embryo/organ clearing, histology, immunochemistry, and imaging

*Whole embryo/organ clearing and imaging.* Embryos and juvenile mice (P0 – P10) were euthanized and hearts and guts were directly dissected. For whole-embryo imaging (Extended Data Figs. 1 and 3a-d), the entire embryo was collected and processed. Adult mice (Fig. 1i, Extended Data Fig. 2) were anesthetized with isoflurane and transcardially perfused with 15 ml cold PBS (pH 7.4) containing 10 U/ml of heparin (Sigma-Aldrich), followed by 25 ml cold 4% PFA before tissue dissection. For some samples (Fig. 1b-c, Extended Data Fig. 2), heart atria were further dissected. The tissues were fixed in 4% PFA at 4 °C overnight and kept in cold PBS at 4 °C before clearing. Tissues were cleared with the CUBIC^60^ method and stained with the following protocol. In brief, dissected tissues were immersed into 1/2-water-diluted reagent-1 (25 wt% urea, 25 wt% Quadrol, 15 wt% Triton X-100) with shaking at 37 °C for 3–6 hours, followed by reagent-1 (R1) with shaking at 37 °C until the tissues are optically cleared. R1 was replaced fresh every two days. Next, the tissue was washed (3× PBS/0.01% NaN3), blocked (2% normal donkey serum, 0.1% Triton X-100, PBS/0.01% NaN_3_) and incubated with indicated primary antibodies (see Supplemental Table 1) in blocking buffer with shaking for 1-2 days at room temperature. Samples were then washed (0.1% Triton X-100, PBS/0.01% NaN_3_) and incubated with fluorophore-conjugated secondary antibodies (see Supplemental Table 1) diluted in blocking buffer with shaking for 1-2 days at room temperature. After antibody incubation, the samples were washed and embedded with 5% low-melting point agarose in 1x PBS. The embedded samples were immersed in 1/2-PBS-diluted reagent-2 (25 wt% urea, 50 wt% sucrose, 10 wt% triethanolamine) overnight at room temperature, and then reagent-2 (R2) at room temperature for 2 days. The samples were finally immersed with mineral oil (Sigma-Aldrich) for at least 1 hour and imaged with the Leica SP8 confocal microscope with a 16× immersion objective (HC FLUOTAR L 16×/0.8 IMM motCORR VISIR, working distance: 8 mm). Heart atria samples (Fig. 1b-c, Extended Data Fig. 2) were immersed in oil and flattened to approximately 500 μm in a custom-built imaging chamber and imaged using the Leica SP8 confocal microscope as described above, with a 10× objective (HC PL APO 10×/0.40 CS2, working distance: 2.1 mm) or a 40× objective (HC PL FLUOTAR L 40×/0.60 CORR, working distance: 3.3 mm). The adult Chat^tdT^ heart sample (Fig. 1i) was imaged with a LaVision Vltramicroscope II light-sheet microscope at the CNNR Imaging Core at Yale University.

*BrdU assay.* BrdU at 100 mg/kg was intraperitoneally injected to pregnant Wnt1^tdT^ female mice (Extended Data Figs. 5d and 6a). After one hour the pregnant mouse was euthanized and embryos were collected. Hearts were dissected and fixed in 4% ice-cold PFA overnight at 4 °C. The samples were cryoprotected in 30% sucrose PBS solution for two days at 4 °C, frozen in OCT, and then stored at −80 °C until cryosection. Cryosections (10 μm) were cut using a cryostat (Thermo Fisher Scientific) and mounted onto Superfrost Plus slides (Thermo Fisher Scientific). Sections were incubated with 1 M HCl for an hour, neutralized with 0.1 M Sodium Borate, pH 8.5 for 10 minutes, washed (3x PBS), permeabilized (0.1% Triton X-100, PBS), blocked (5% Normal Donkey Serum, PBST (PBS, 0.1% Triton X-100)) and incubated with primary antibodies (see Supplemental Table 1) diluted in blocking buffer at 4 °C overnight. Then, the slides were washed (3x PBST), and incubated with fluorophore-conjugated secondary antibodies (see Supplemental Table 1) diluted in blocking buffer for 2 hours at room temperature. After incubation, the samples were washed (3x PBST), and mounted with Prolong^TM^ Diamond Antifade Moutant with DAPI (Thermo Fisher Scientific) prior to imaging with the Leica SP8 confocal microscope.

### Geometric transformation of 3D heart images

Locations of PHOX2B^+^ ICNS nuclei and heart tissue outlines were measured using Fiji and imported into MATLAB for geometric transformation (Extended Data Fig. 4a). Four landmarks (anterior, posterior, right, and left) on the heart atrium that encompass the ICNS and define the ICNS plane (Extended Data Fig. 4b) were selected for each sample to transform all imaged hearts into the same orientation and position. The ICNS reference plane was defined as the plane with the least cumulative Euclidean distance to all four landmark points and its normal vector was calculated (Extended Data Fig. 4c). As anterior and posterior landmarks were selected on the midline of the heart atrium, their midpoint was used as the center of the reference plane, and a second reference plane that contains both anterior and posterior landmarks and is perpendicular to the ICNS reference plane was calculated. Then a third reference plane was constructed to complete the three new orthogonal axes of the heart sample after transformation. Transformed coordinates of ICNS cells were then calculated using the new axes (Extended Data Fig. 4d). As ICNS cells are predominantly distributed on the ICNS reference plane, their z-axes are negligible after transformation and thus can be compared using a 2D plot (Extended Data Fig. 4d).

We subsequently delineated the transformed 2D ICNS reference plane, partitioning it into 50 μm x 50 μm grids. We converted the 2D ICNS reference plane into a binary image and applied the following methods to compute the ICNS features described in Fig.1f and Extended Data Fig. 4. A grid containing more than two ICNS cells was counted as an ICNS-containing grid. The ICNS cluster area was calculated as the total area of ICNS-containing grids (number of ICNS-containing grid x grid area). Density is calculated by the total number of ICNS cells in clusters divided by the ICNS cluster area. To determine the cluster number, we first calculated the highest ICNS number per grid within the sample (max^ICNS^). Subsequently, we defined boundry of individual ICNS clusters as the grids with ICNS cells less than 20% of max^ICNS^. Individual ICNS clusters were determined using the following criteria: (1) it has clear boundaries, non-overlapping with other clusters, and (2) its peak density (highest ICNS number per grid) is more than 50% of max^ICNS^. To determine the distribution variability, we first normalized the 2D ICNS reference plane area to the distance between anterior and posterior landmarks for all samples. We then re-partitioned the normalized 2D ICNS reference plane into 0.1 x 0.1 grids. Next, we calculated the probability of ICNS distribution for each grid by dividing the number of ICNS cells within the target grid by the total number of ICNS. Finally, we calculated for each sample the distribution variability across all grids. The distance to the nearest ICNS cells was determined based on the distance to the ten closest ICNS cells. The inner area was delineated by encircling the region between ICNS clusters using Fiji.

### Simulation of ICNS organization

The organization of ICNS cells during proliferation of surrounding heart cells in Phase II was simulated using MATLAB (Fig. 1h, Supplementary Video 1). Three groups of cells were defined to mimic the distribution of cardiac and ICNS cells: G1 (green) represents the cells in the “inner area” of the ICNS ganglion, G2 (red) represents ICNS cells, and G3 (blue) represents the outer cardiac cells. The initial cell density of the three groups was set the same. The proliferation rates of G1 and G3 were set at 0.5, while the proliferation rate of G2 was set at 0.01, recapitulating our observation as shown in Extended Data Fig. 5d. For each proliferation round, the number of proliferating cells in each cell group was calculated as the total number of cells in that group at the time times its proliferation rate. Proliferating cells were randomly selected within the group, and two daughter cells were generated from the selected cell at the same spatial location. Cells were then allowed to freely migrate along the cell density gradient to less occupied locations. In addition, once isolated, G2 cells were set to move towards the center of their 10 closest G2 neighbour cells to mimic the clustering of ICNS cells. Each proliferation round contained a certain number of migration steps, followed by the next proliferation round. The distribution of all cells, colored by their identities, was saved and plotted after each simulation step, which together constitutes the stimulation pseudotime.

### Label Free Quantitative mass spectrometry analysis

E14.5 hearts from Wnt1^L10-GFP^ mice were dissected on ice under a Leica M205FCA Fluorescence Stereo Microscope to separate the atrium region that contains fluorescently labelled ICNS cells (heart^near^) and the atrium region that doesn’t (heart^far^). Dissected tissues were collected into 1.5-mL tubes, snap frozen with liquid nitrogen, and stored at -80 °C. Each sample contains approximately 1 mm^3^ tissues from four embryos. Sample were subjected to Labelled Free Quantitative Mass Spectrometry analysis at the Keck Biotechnology Resource Laboratory at Yale University.

### Coculture ENS cells with organ cells

#### Cell dissociation and culturing

The heart and gut were dissected from E13.5 Phox2b^tdT^ embryos on ice under a Leica M205FCA Fluorescence Stereo Microscope. Heart atria containing the ICNS were collected. The mesentery of the gut was removed, and the entire intestine were used for ENS cell dissociation. Cell dissociation was performed as described in the ‘**Single-cell RNA-sequencing analysis**’ section. To prevent contamination, 100 U/mL penicillin, 100 μg/mL streptomycin, and 100 μg/mL Gentamicin were supplemented throughout the dissociation process. Dissociated heart and gut cells were subjected to FACS using a BD FACS Aria cell sorter at the Yale Flow Cytometry Facility. tdTomato positive and negative cells were collected into separate tubes containing ice-cold 10% L-15 media. Cells were centrifuged at 700 x g at 4 °C for 10 minutes, resuspended in culture media (Neurobasal^TM^ Plus Medium (A3582901, Gibco) supplemented with 1% FBS, 2 mM L-Glutamine, B-27^TM^ Plus Supplement (A3582801, Gibco), 100 U/mL penicillin, 100 μg/mL streptomycin, and 100 μg/mL Gentamicin), and plated onto 24-well cell culture plates (Corning, 3,000 tdTomato^+^ ENS cells with 6,000 tdTomato^-^ gut or heart cells, mixed) pre-coated with a thin layer of Collagen I/Matrigel (#356237, Corning) mixture that contains 0.5 mg /mL of Matrigel and 33 μg/mL Collagen I (ALX-522-440-0050, Enzo Life Sciences). Cells were then incubated at 37 °C in 5% CO_2_ and co-cultured for 4-5 days.

#### Single-cell RNA-sequencing of cocultured cells

After removing cell culture media, 2 mg/mL Collagenase A and 2 mg/mL of Dispase II solutions were added to the cells and incubated for 10 minutes at 37 °C. The cell solutions were collected into 10% FBS L-15 media on ice. This step was repeated 3-6 times until the coating layer was completely digested. Cells were centrifuged at 200 x g for 10 minutes at 4 °C, washed with fresh 10% FBS L-15 media, and incubated with Corning Cell Recovery solution (#354253) on ice for 20 minutes to dissolve any residual Matrigel. After washes, cells were manually triturated with a Pasteur pipet, strained through a 40 μm cell strainer, washed, and resuspended in 1x HBSS, 0.04% BSA for single-cell RNA-sequencing as described in the ‘**Single-cell RNA-sequencing analysis**’ section.

